# Competitive integration of time and reward explains value-sensitive foraging decisions and frontal cortex ramping dynamics

**DOI:** 10.1101/2023.09.05.556267

**Authors:** Michael Bukwich, Malcolm G. Campbell, David Zoltowski, Lyle Kingsbury, Momchil S. Tomov, Joshua Stern, HyungGoo R. Kim, Jan Drugowitsch, Scott W. Linderman, Naoshige Uchida

## Abstract

Patch foraging presents a ubiquitous decision-making process in which animals decide when to abandon a resource patch of diminishing value to pursue an alternative. We developed a virtual foraging task in which mouse behavior varied systematically with patch value. Mouse behavior could be explained by a model integrating time and rewards antagonistically, scaled by a latent patience state. The model accounted for deviations from predictions of optimal foraging theory. Neural recordings throughout frontal areas revealed encoding of decision variables from the integrator model, most robustly in frontal cortex. Regression modeling followed by unsupervised clustering identified a subset of ramping neurons. These neurons’ firing rates ramped up gradually (up to tens of seconds), were inhibited by rewards, and were better described as a continuous ramp than a discrete stepping process. Together, these results identify integration via frontal cortex ramping dynamics as a candidate mechanism for solving patch foraging problems.

## Introduction

One of the most fundamental decisions that animals must regularly make is how to effectively allocate time in the pursuit of resources. Patch foraging—the process of deciding when to leave a location with depleting resources (e.g. food) to search elsewhere—is a classic problem of this type that has been studied in the framework of optimal foraging theory in behavioral ecology^1,2^. Theoretical work has shown that foraging animals should leave a patch when the instantaneous rate of resource intake drops to the average rate in the environment—a result known as the Marginal Value Theorem (MVT)^3^. Although this theorem provides a mathematical solution to optimal patch-leaving decisions, (1) whether the behavior of animals conforms precisely to the MVT remains debated, and (2) the biological processes underlying patch-leaving decisions remain to be determined^1,4-6^.

The MVT defines an abstract decision rule but does not explain the underlying decision process.

Efforts have therefore been made to identify simple patch-leaving rules that can approximate optimal decisions. One such rule is a “giving-up time rule”: leaving after a predetermined amount of time has elapsed since the last incident of resource intake^7-9^. Another class of decision rule is formulated as an integrate-to-threshold process, similar to those accounting for perceptual decisions studied in psychology and neurobiology, often structured as drift-diffusion models^10-15^. In the context of patch foraging, these models track a decision variable (DV) which integrates elapsed time and intermittent occurrence of resource intake in opposite directions and trigger patch leaving when the DV reaches a threshold level.

Because of the competitive interactions between elapsed time and resource intake, a patch-leaving decision is naturally delayed within patches containing more resources. This type of model was first applied in the context of patch-foraging to egg-laying behavior in the parasitoid wasp, *Nemeritis canescens*^*16*^. Furthermore, recent theoretical work has demonstrated that integrate-to-threshold models can provide MVT-optimal decisions under idealized conditions^17^. This study further showed that integrator models can achieve near-optimal behavior in more realistic conditions and, through parameter tuning, can also implement non-MVT strategies, such as counting, which can be optimal when assumptions of MVT are violated. The proposition that integrator models can be adapted to solve foraging problems is attractive because they offer a potential mechanism through which to link foraging behavior to its neural basis.

Numerous studies have demonstrated the correspondence between frontal cortex activity and foraging decisions^18^. Hayden et al. (2011) modeled patch-foraging using an eye movement-based task in monkeys and showed that neurons in the anterior cingulate cortex increase their pre-saccadic transient responses up to a certain level before leaving a patch^19^. Work in humans has similarly implicated regions in frontal cortex in tracking value signals during foraging-like decisions^20,21^. Studies of when-to-leave decisions have identified multiple areas of rodent frontal cortex^22,23^, including anterior cingulate cortex (ACC)^24^, orbitofrontal cortex (OFC)^25,26^, and secondary motor cortex (M2)^23,27^ as playing a causal role in leaving decisions. However, the neural basis of the decision process remains to be identified.

In this study, we set out to identify the behavioral algorithms and neural activity patterns that support a classic patch leaving problem. We first present a novel head-fixed virtual reality patch foraging task for mice. Our results show that mouse behavior matched the MVT qualitatively, yielding greater patch residence time (PRT) on patches in which rewards were more abundant. However, stronger tests for the MVT, exploiting our task designs, showed that the behavior systematically deviated from MVT predictions. We show that, instead, integration-to-threshold models that were globally scaled by a slowly varying “patience” variable estimated from surrounding trials can explain not only the overall behavioral patterns, but also these systematic deviations with quantitative precision. Furthermore, we observed a prevalence of neurons which exhibited a slow, ramp-to-threshold type activity that was decremented by intermittent rewards. Dynamically updating decision variables obtained from integration models could be predicted from neural activity in the frontal cortex, and vice versa, more accurately than in subcortical areas. These results suggest that mice could solve the patch foraging problem via an integration process within the frontal cortex.

## Results

### A head-fixed patch foraging task for mice

We developed a patch foraging task for head-fixed mice in virtual reality (Fig. 1), in which they ran on a cylindrical treadmill to traverse a linear corridor. After mice ran a fixed distance, a visual “proximity cue” was presented, indicating the availability of a resource patch. If mice stopped running while the proximity cue was on, a patch trial was initiated with a visual cue, and water droplet rewards were delivered stochastically while mice remained on the patch. Each patch delivered water droplets of one of three sizes (1, 2, or 4 *μL*), which was consistent within a patch. Rewards were delivered probabilistically each second on the patch, with one of three initial reward probabilities (0.125, 0.25, or 0.5) followed by exponential decay of that probability with time constant *τ* = 8 *sec*. There were thus nine patch types: three reward sizes × three reward frequencies (Fig. 1B). To encourage stopping on patches, the first reward was delivered deterministically on all trials (at *t* = 0, the time of stop). Mice could then choose to remain on the patch for any amount of additional time, during which additional probabilistic reward deliveries might occur at one-second increments. Mice were free to leave a patch any time by travelling a fixed distance on the wheel, at which point they would enter the inter-patch-interval and be required to run to the next patch to receive more water.

**Figure 1:**
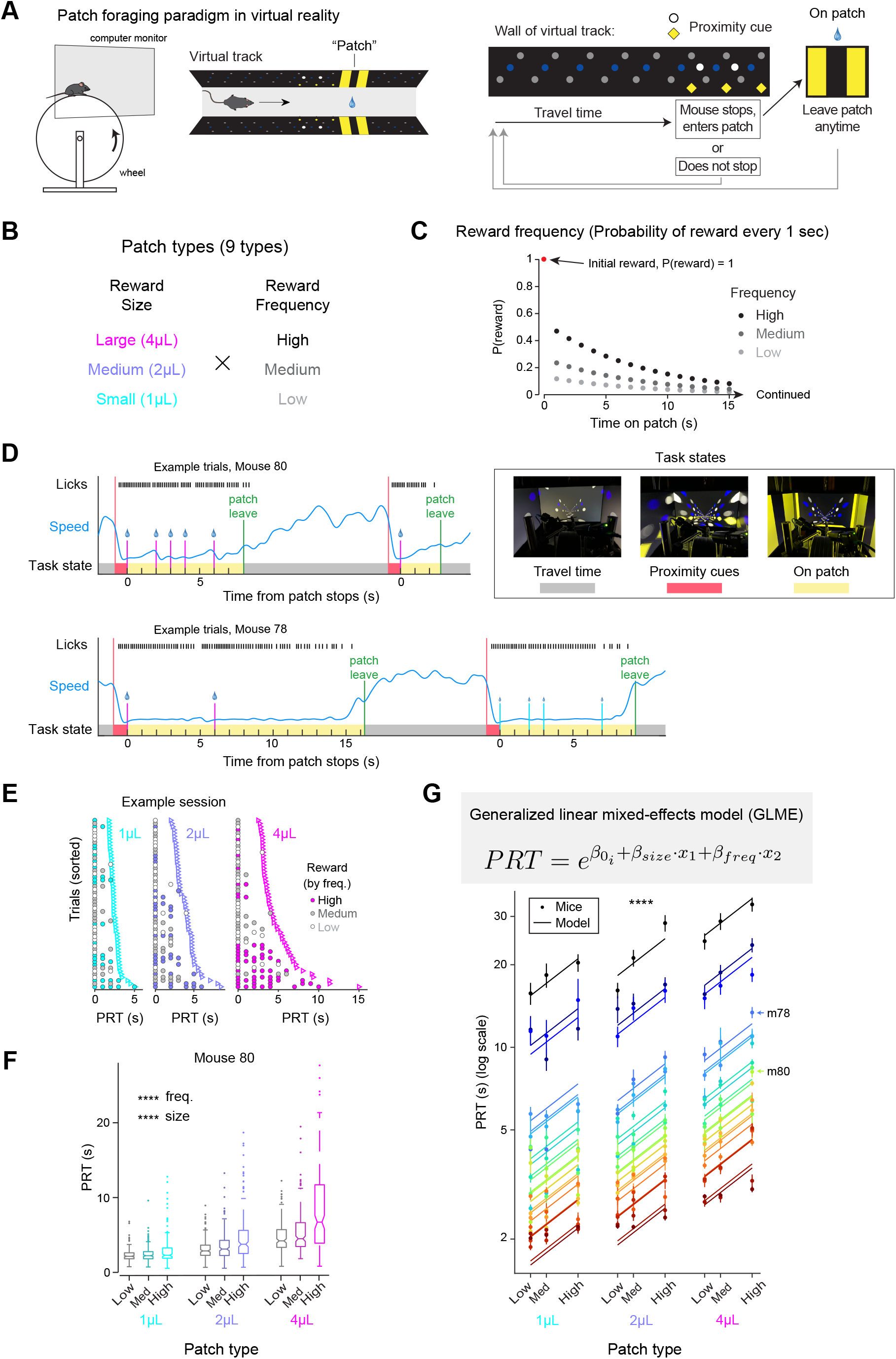
Mice foraging times calibrate to reward statistics. A) Virtual linear track for patch foraging task. B) Combinations of three reward sizes and frequencies yield nine patch types. C) Probability of reward delivery after each one-second interval per frequency condition. D) Example trials from two mice. Mice stop running in response to the proximity cue to enter patches and receive stochastically delivered water rewards. Photographs of the three task states are shown. Monitor brightness was increased for the photos. E) Reward deliveries and patch leave times from an example session grouped by patch reward size. Trials are sorted in ascending order of PRT from top to bottom per reward size. Dots are shaded based on underlying reward frequency condition. F) PRT per patch type from an example mouse. Column groupings separate patches by reward size. Within columns, patches are split by reward frequency. G) A generalized linear mixed-effects model (GLME) with log link function predicts mice’s PRTs across conditions. To demonstrate the log-linearity of the fit, frequency conditions are spaced proportionally to their values (0.125:0.25:0.5). Dots indicate empirical mouse PRT with error bars showing standard error of the mean. Lines represent GLME fits. Colored by mouse ID (in GLME equation, ‘i’ = mouse ID).

### Patch residence times are a scaled function of reward statistics

During the foraging task, mice consistently guided their behavior towards obtaining water rewards. Example trials from two mice demonstrate how their running and licking evolved in response to task events (Fig. 1D), stopping in response to proximity cues and licking in expectation of rewards.

Reward-seeking behavior was evident across the population. In response to the proximity cue, mice successfully stopped to enter most patches (Fig. S1A). As mice stopped to enter a patch, they showed anticipatory licking at the same time as they began decreasing their speed (Fig. S1B), demonstrating an expectation of reward availability. After receiving an initial reward at *t* = 0 (consistent across all patch trials), they began modulating their behavior in response to reward size. Lick rates were higher on patches with larger reward size, and mice waited longer before increasing their running speed (Fig. S1B).

The canonical signature of value-sensitivity during patch foraging is the dependence of patch residence time (PRT) on resource richness. Mice indeed showed higher PRT in patches with larger and/or more frequent rewards (example session, Fig. 1E; example mouse, Fig. 1F; all mice, Fig. 1G), though with marked differences in overall willingness to wait across subjects. In contrast to the broad differences in mean PRT across subjects, we observed that relative PRTs across patch types were approximately co-linear across mice after log-scaling (Fig. 1G). This trend occurs because the influence of reward statistics on waiting time scales systematically with mean PRT (Fig. S2A). We therefore tested whether a generalized linear mixed-effects model (GLME) using a log link function could capture variability of mean PRT across mice and patch types. We fit a 3-factor GLME with reward size and frequency as fixed effects and mouse identity as a random effect and found that this model captured mean PRTs across conditions for the population (*β*_*size*_ = 0.168, *β*_*freq*_ = 0.103, both p < 0.0001, Fig. 1G).

In addition to the differences in waiting behavior across mice, we observed substantial within-subject variability in PRT over time. Some mice displayed striking fluctuations in their PRTs across sessions (Fig. 2A, Left: example mouse. Fig. S1E, population). PRT also varied gradually within sessions such that PRTs on successive patch trials were correlated with one another (Fig. 2B). This suggests a slowly fluctuating latent state, which we henceforth call ‘patience’, that describes the varying mean levels of willingness-to-wait we observed in mice. Taking advantage of this, we used PRT from successive patches as a proxy for estimating subjects’ latent degree of ‘patience’ on a given trial (Fig. 2C, Left). We then asked whether using these latent patient estimates to scale a common function could account for variance in PRTs across the population. We modified our GLME, transforming it into a GLM by replacing the random effect term for mouse identity with a regressor for latent patience estimates (Fig. 2D). Using a single set of four free parameters shared across mice, GLM-fitted PRTs well matched mice’s mean PRTs across patch types (Fig. 2D) and robustly predicted single trial PRTs for most mice (Population, median R^2^ = 0.45, Fig. 2E; example mouse, Fig. S2G, Left).

**Figure 2:**
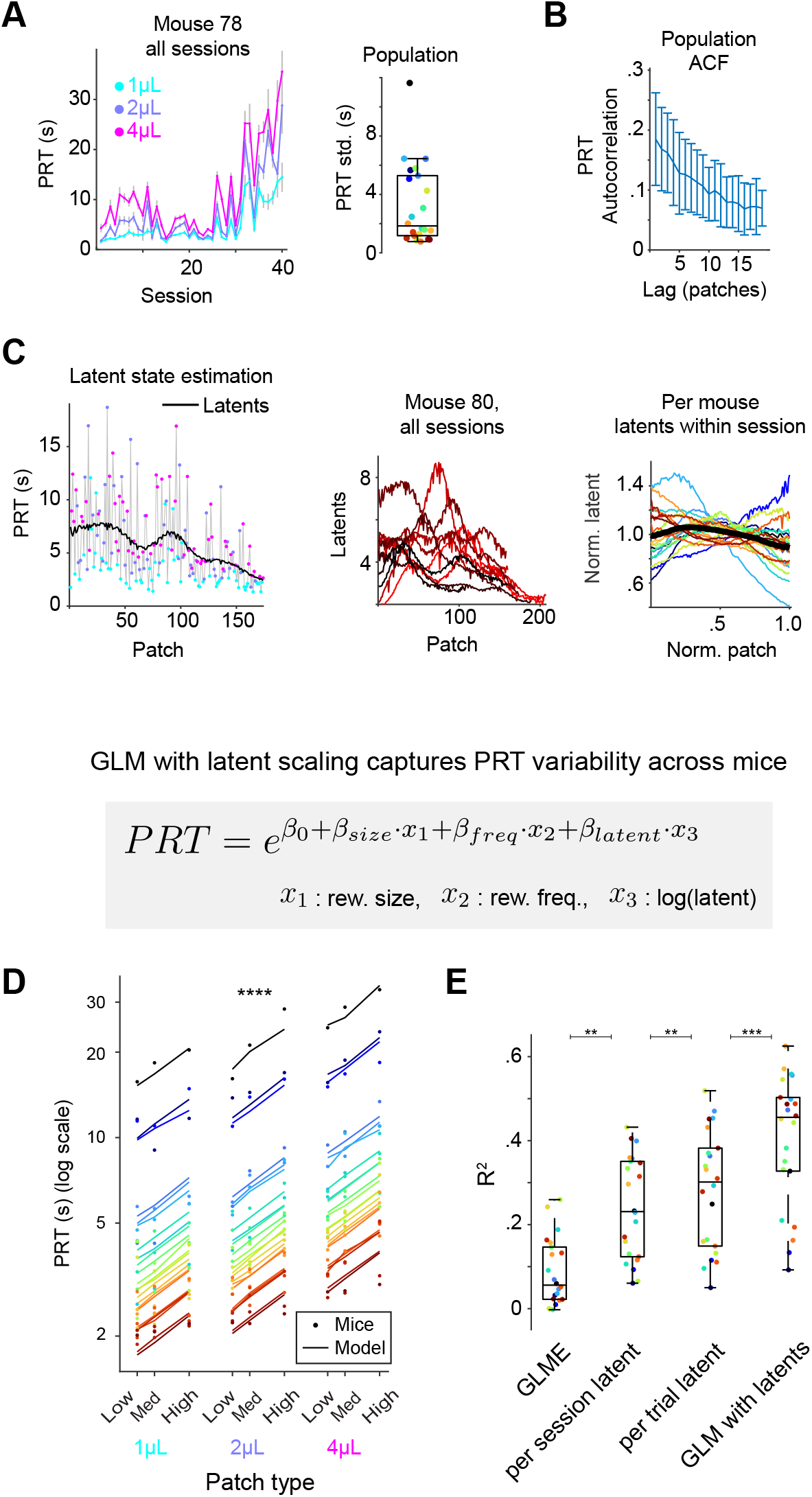
PRT variance is explained by scaling a common function. A) Left: Mean PRT per μ*L* across sessions, example mouse. Right: Standard deviation of mean PRT across sessions per subject, colored by mouse ID. B) Autocorrelation function of PRT over patches, mean across subjects. Error bars indicate standard deviation across subjects. C) Applying a Gaussian filter of PRT over successive trials provides an estimate of latent state. Left: Example session latent state inference (mouse 80). Gray trace tracks PRT across patches with colored dots indicating reward size for each patch. Black trace indicates value of the latent state estimation. Middle: Estimated latent across trials over all 10 sessions from mouse 80. Right: Mean normalized latent estimates across sessions per mouse. D) A generalized linear model (GLM) scaled by latent patience estimates accounts for PRTs across mice. Dots indicate empirical mouse PRT. Lines represent GLM fits. E) R^2^ statistics for predicting PRT using the GLME (from Fig. 1G), mean latent per session, per trial latent, and the latent-scaled GLM. Models ordered by ascending levels of R^2^, stars indicate significance of pairwise Wilcoxon signed-rank tests (p = .0044, .0027, .002, Bonferroni-adjusted).

### Mice behavior deviates from MVT predictions

We next performed direct tests of MVT predictions. First, the MVT dictates that, if patches are monotonically depleting in value as resources are consumed, an optimal forager should leave a patch once the instantaneous rate of reward drops to the average expected reward rate of the environment. Thus, irrespective of initial resource abundance, patches should all be left around this same threshold crossing (Fig. 3A, Left; **Prediction 1**, see Methods). To test this prediction of MVT, we computed expected reward rate at the time of each patch leave. We found that mouse behavior strongly violated this prediction, with animals leaving high value patches too early and low value patches too late^28^, relative to an ideal observer, with the dominant effect driven by reward size (Fig. 3A, middle: example mouse, p < 0.0001 for reward size and p > 0.99 for reward frequency, 2-way ANOVA, Bonferroni-adjusted; right: population means, p < 0.0001 for both size and frequency, linear mixed-effects model).

**Figure 3:**
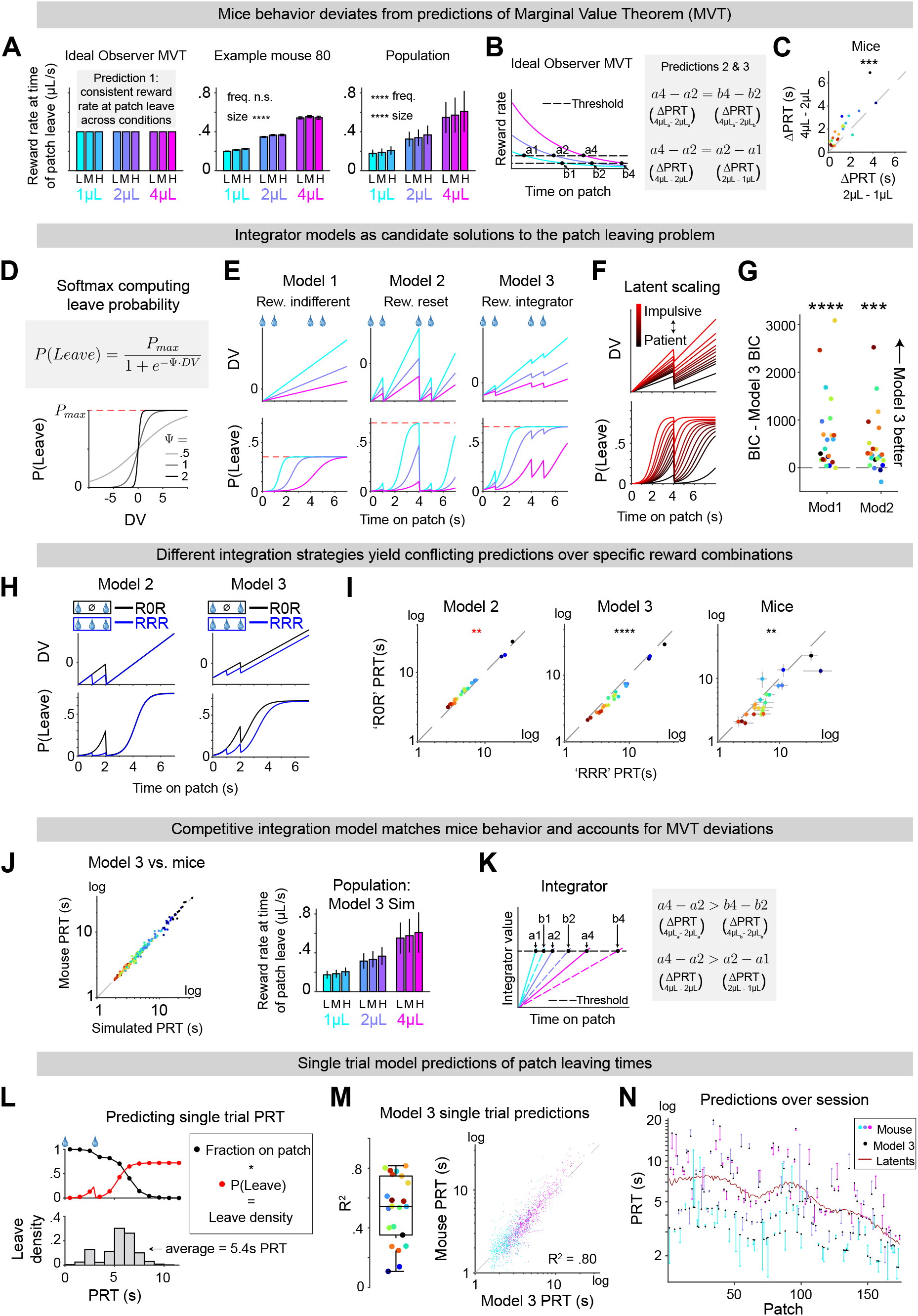
A competitive integration process explains foraging behavior. A) Instantaneous expected reward rate at time of patch leave across patch types, predicted by optimal MVT (Left), sample mouse (Middle, Bonferroni-adjusted p < 0.0001 for size, p > .99 for frequency, 2-way ANOVA), and population (Right, p < 0.0001 for size and frequency, linear mixed-effects model). Error bars for sample mouse indicate standard error of the mean. Error bars for population average indicate standard deviation across mice. B) Schematic demonstrating MVT predictions for PRT per reward size for two different thresholds. Traces are colored by reward size. Dashed lines show two sample thresholds, *a* and *b*. Black dots indicate points of threshold crossings for each reward size. Gray box notes examples for **Prediction 2** (Top) and **Prediction 3** (Bottom). C) Mice show greater differences in mean PRT between 4μ*L* and 2μ*L* patches, compared to 2μ*L* and 1μ*L* patches. D) Sigmoid transformation of decision variable (DV) into probability of patch leave per one-second interval, scaled by different inverse temperature values (light grey = 0.5, dark grey = 1.0, black = 2.0). Red dashed line indicates the maximum *P*(*Leave*) output, *P*_*max*_. E) Example schematics for DV (Top) and corresponding *P*(*Leave*) output (Bottom) for three different integrator models over patches with rewards delivered at *t* = [0,1,4,5] seconds. DVs ramp upwards over time. Model 1 does not respond to reward deliveries. Model 2 resets to its baseline value following any reward delivery. Model 3 integrates rewards with a constant negative value. Red dashed line indicates the maximum *P*(*Leave*) output, *P*_*max*_. F) Schematic demonstrating how *DV* and *P*(*Leave*) scale relative to latent patience estimation across patches, for a sample patch in which rewards were delivered 0 and 4 seconds. Black traces indicate patches with more patient estimations and ramp up more slowly. Red traces indicate impulsive latents and yield sharper ramps. G) Relative BIC values for the model fits across subjects. Model 3 yields a superior fit to Models 1 and 2 (p < 0.0001, p = 0.0009, Wilcoxon signed-rank test). H) Example schematics demonstrating how Models 2 (Left) and 3 (Right) make differing predictions for DV (Top) and *P*(*Leave*) predictions (Bottom) on patches with rewards at *t* = [0,2] sec (‘R0R’ patches, black) compared with patches with rewards at *t* = [0,1,2] sec (‘RRR’ patches, blue). I) Per subject mean Simulated PRT for ‘R0R’ versus ‘RRR’ patches from Model 2 fits (Left), Model 3 fits (Middle), and empirical mice PRT (Right). Model 3 and mice PRTs were significantly higher for ‘RRR’ versus ‘R0R’ trials (p < 0.0001, Model 3; p = 0.0024, Mice; Wilcoxon signed-rank test). There was a small but significant effect of greater PRTs for ‘R0R’ versus ‘RRR’ trials for Model 2 (p = 0.0037). However, this resulted due to a selection bias over latent patience values and was in the opposite direction of the empirical mice data. Points are colored per mouse. Axes are log-scaled. J) Mean PRT across patch types, per subject from Model 3 simulations versus empirical mouse PRT, log-scaled (*R*^2^ = 0.985, over patch type means, Left), points colored per mouse. Instantaneous expected reward rate at patch leave across patch types from population simulations of Model 3 using fit parameters per mouse. K) Schematic demonstrating how integrator models can account for deviations from MVT predictions. Example traces are shown when patience is lower (solid lines) versus higher (dashed lines) for each reward size (colored per µL). Patch leave times are determined by integrator value reaching threshold (black dashed line, dots indicate threshold crossings). Gray box notes example violations of **Prediction 2** (Top) and **Prediction 3** (Bottom). L) Schematic depicting a sample trial for calculating model-predicted PRT on single trials. Black dots indicate fraction of trials the model predicts subjects would still be on the patch at the start of that one-second time bin (Top). Red dots indicate the probability of leaving, *P*(*Leave*), within that time bin. The product of fraction of trials remaining on patch times and *P*(*Leave*) determines the corresponding leave density for that time bin (Bottom). The average predicted PRT is then calculated as that trial’s prediction. M) *R*^2^ statistics for single trial predictions from cross-validated Model 3 fits across mice (Left, median *R*^2^ = 0.544). Box edges indicate 25^th^ and 75^th^ percentiles. Whiskers stretch out to most extreme points, as none were categorized as outliers. Example mouse single trial Model 3 predicted PRT versus empirical mouse PRT, points colored reward size per patch (Right, R^2^ = 0.801). Predicted PRT for each trial was calculated using model fit parameters from training folds and compared with PRT over trials from held-out test folds. N) Model 3 predicted PRT and empirical mouse PRT across patches from the example session in left panel of Fig. 2C. Colored dots indicate mouse PRT, colored by reward size. Black dots indicate Model 3 predicted PRT, calculated from fitting on training folds. Lines connecting dots highlight the difference in predicted versus empirical PRT per trial. Red trace indicates latent estimations of patience for each patch. Y-axis is log-scaled.

Second, we considered more specific predictions of MVT that can be tested because of our task’s reward schedules. Because all patch types had exponentially decaying reward probabilities with the same time constant (Fig. 1C), differences in PRTs across patch types should remain constant, regardless of the threshold (Fig. 3B; **Prediction 2**, see Methods). Furthermore, because we used patches with proportional reward sizes (1, 2, and 4μ*L*), MVT predicts that the difference in leave time between 4 and 2 *μL* patches (*PRT*_4μ*L*_ − *PRT*_2μ*L*_) should be equal to the difference between 2 and 1 *μL* patches (*PRT*_2μ*L*_ − *PRT*_1μ*L*_; Fig. 3B; **Prediction 3**, see Methods). Neither of these predictions was borne out in the data. Instead, reward sensitivity grew with mean PRT (Fig. 2D, Fig. S2) and we consistently observed *PRT*_4μ*L*_ − *PRT*_2μ*L*_ > *PRT*_2μ*L*_ − *PRT*_1μ*L*_ (Fig. 3C, p = 0.0002, Wilcoxon signed-rank test).

### A competitive integration process explains patch leaving decisions

Prior work has demonstrated the viability of integrator models as a potential solution to the patch leaving problem, as they can achieve MVT optimal behavior in limiting cases and produce satisfactory performance under more naturalistic conditions^17^. To test whether integrator models can match the observed PRTs of foraging mice, we devised several integrator models which differentially track time and reward events, yielding probabilistic predictions of patch leaving decisions. For each model, we compute a decision variable, *DV*, for each time bin, which is then transformed through a softmax function to generate a predicted probability, *P*(*Leave*), of the animal leaving the patch—i.e., the hazard rate of leaving (Fig. 3D; see Methods for detailed model description). As *DV* increases, so does *P*(*Leave*).

Because the model must account for decision processes unfolding over time durations spanning orders of magnitude, we constrained the maximum probability to *P*_*max*_, which is a free parameter. As a result, the midpoint of the sigmoid, corresponding to instances when the *DV* = 0, is not an indifference point wherein *P*(*Leave*) = 0.5, but instead is the point at which 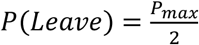.

We fit three distinct integrator models for computing DVs over time (Fig. 3E). In each model, a *DV* ramps up linearly over time in the absence of rewards, with its slope scaled by reward size via a power law function. What distinguishes the models is how they respond to reward deliveries. Model 1 exclusively tracks time on patch, and is indifferent to rewards, ramping upwards irrespective of additional reward deliveries. Model 2 is a full reset model, which returns to its initial baseline value upon each reward delivery. Model 2 is therefore “memoryless,” in the sense that its value is dictated only by the time since the most recent reward and the patch’s reward size. It is equivalent to the “giving-up time model” discussed earlier^7^, with a giving-up time defined for each reward amount. Model 3 performs a competitive integration process of rewards versus time, preserving effects of multiple rewards over time instead of only the most recent one. For each reward, Model 3 is reduced by an amount governed by a free parameter, *R*. For the purposes of this model, we fit only a single value for *R* rather than scaling it with reward size, as we found that scaling slope alone was sufficient to account for the observed differences in PRT with reward size. As mouse PRTs and reward sensitivity were both strongly influenced by latent levels of patience (Fig. 2, S2), we scaled model parameters by the latent state on each trial (Fig. S3A; Methods). Non-latent-scaled versions of each model performed substantially worse than their latent-scaled counterparts, with scaling of slope and *P*_*max*_ consistently yielding the best fits (Fig. S3A). We therefore used this scaling for our models (Fig. 3F).

To better understand how rewards affect the decision process, we compared predicted leave probabilities from the three integrator models to observed patch-leaving behavior. Model 1 can be ruled out a priori, as it could not account for the observed dependency of PRT on reward frequency, but we included it still as a baseline model for comparison. In contrast, Models 2 and 3 could both potentially account for the reward effects qualitatively, as each predict larger PRT for higher reward size and frequency by reducing their *DV* and *P*(*Leave*) predictions following reward deliveries. We therefore sought a systematic means of comparing model performance beyond predicting mean PRT, instead focusing on the probabilistic leave decisions the models predicted within individual trials.

To fit and compare the models, we fit their parameters for each mouse separately by maximum likelihood and compare their goodness of fit using the Bayesian Information Criterion (BIC) (Fig. 3G), to account for the different number of parameters across models. BIC was lowest for Model 3, demonstrating that the competitive integration model best explained mouse leaving behavior across trials (p < 0.0001, Model 3 over Model 1; p = 0.0009, Model 3 over Model 2; Wilcoxon signed-rank test, Bonferroni correction). We corroborated the BIC metrics using 5-fold cross-validation and obtained similar results, with the likelihood consistently higher for Model 3 than the alternative models (Fig. S3B, p < 0.0001, Model 3 over Model 1; p = 0.0008, Model 3 over 2, Wilcoxon signed-rank test, Bonferroni correction).

To further distinguish between Models 2 and 3, we took advantage of the stochasticity and the richness in trial history that our task provides and examined trial types with specific reward histories for which Models 2 and 3 make divergent predictions (Fig. 3H). We compared two combinations of reward sequences: ‘RRR’ trials, which delivered rewards at *t* = 0,1,2 seconds, and ‘R0R’ trials, which delivered rewards *t* = 0 and 2 seconds (but no reward at *t* = 1). Because Model 2 produces a full reset following reward, it predicts identical leaving behavior on both trial types, as both reset to the same value after the final reward (Fig. 3H, left). In contrast, Model 3 integrates reward history and preserves the effect of the *t* = 1 reward, predicting higher PRT on ‘RRR’ over ‘R0R’ patches (Fig. 3H, right). Simulations demonstrate these differences (Fig. 3I, left, middle), and mouse behavior matched Model 3 predictions, with nearly all subjects waiting longer on ‘RRR’ trials (Fig. 3I, right). These results support Model 3’s competitive integration over Model 2’s reward reset (a.k.a. “giving-up time” model) as a better description of the decision algorithm. We similarly compared ‘RR0’ versus ‘R0R’ trials and found that mouse behavior again better matched Model 3 predictions (Fig. S3C-D).

Using our Model 3 parameter fits for each mouse, we simulated PRTs by using *P*(*Leave*) outputs to stochastically generate patch leaving decisions. Mean simulated PRTs per patch type from Model 3 closely matched mice’s empirical PRTs across the population (Fig. 3J, Left, MSE = 0.413s). Moreover, we performed the same calculation of instantaneous expected reward rate at the time of patch leave (from Fig. 3A) on our simulated data and found that Model 3 produced a nearly identical deviation from MVT **Prediction 1** as was observed in our experiments (Fig. 3J, Right).

Figure 3K demonstrates how integrator models can implicitly explain the departures from MVT predictions that we observed in the foraging task, with most parameter combinations resulting in different terminal reward rates across patch types. For example, given two integration processes with different slopes (e.g. 2µL versus 4µL traces shown, solid lines), proportionally reducing their slopes (corresponding to increased patience, dashed lines) results in a greater PRT difference between them, thus violating **Prediction 2**. Additionally, unless slopes are precisely tuned across reward sizes, then *PRT*_4μ*L*_ − *PRT*_2μ*L*_ ≠ *PRT*_2μ*L*_ − *PRT*_1μ*L*_, thereby violating **Prediction 3**. Consistent with our experimental results shown in Fig. 3C, our best-fitting model replicated the observed *PRT*_4μ*L*_ − *PRT*_2μ*L*_ > *PRT*_2μ*L*_ − *PRT*_1μ*L*_.

Notably, prior work has emphasized that integrator models can achieve MVT optimal behavior under idealized conditions^17^; here, we show that while mouse wait times increase with patch value (a core feature of optimal foraging), they *deviate* from optimality in ways that can be explained by a patience-scaled integration process that is not perfectly tuned to reward statistics. Therefore, while we cannot rule out additional assumptions that may render our data consistent with MVT (e.g. imperfect task knowledge or risk sensitivity), our results suggest that integrator models offer a parsimonious explanation for the observed behavior.

We next asked how well we could predict PRT across individual held-out trials not used to fit Model 3 (Fig. 3L, example trial; Methods). We found that model predicted PRT distributions explained a substantial amount of the variance across individual patches (Fig 3M, Left: population, median *R*^2^ = 0.54, Right: example mouse, *R*^2^ = 0.80). An example session is shown to demonstrate how these model predictions capture PRT variance beyond that which is predicted by the latent patience estimates (Fig. 3N, Fig. S3H).

In addition to our per mouse model fits, we fit a version of Model 3 in which a single set of free parameters was shared across mice. With the differences across mice being driven solely by the latent patience scaling, we were able to robustly capture the variability across mice and conditions consistently better than an alternative Model 3 fit separately per mouse without latent scaling (Fig. S3F, population; Fig. S3G, sample mice). These results demonstrate that latent-patience scaling is both necessary and sufficient to account for the variability in PRT that we observe across subjects and patches, providing further evidence that individual differences might be driven via scaling of a common underlying algorithm.

### Slow ramping is a prominent feature of frontal cortex activity

The above behavioral analyses have indicated that various aspects of behavior in our patch foraging task, including systematic deviations from the MVT, can be parsimoniously explained by an integration-to-threshold model. We next aimed to investigate the neural underpinnings of the foraging strategies identified above. We recorded spiking activity of neurons using high density probes, Neuropixels, broadly throughout frontal cortex and underlying subcortical areas while mice performed the task (Fig. 4A,B; n = 6,090 units from 33 recording sessions in 9 mice). We observed neurons whose activity slowly ramped up or down as mice remained in the patch (example in Fig. 4C). For this example neuron, firing rate ramped to a similar level prior to patch leave with a slope that depended on reward size, and was inhibited by reward delivery—much like the ramping DVs in our behavioral model.

**Figure 4:**
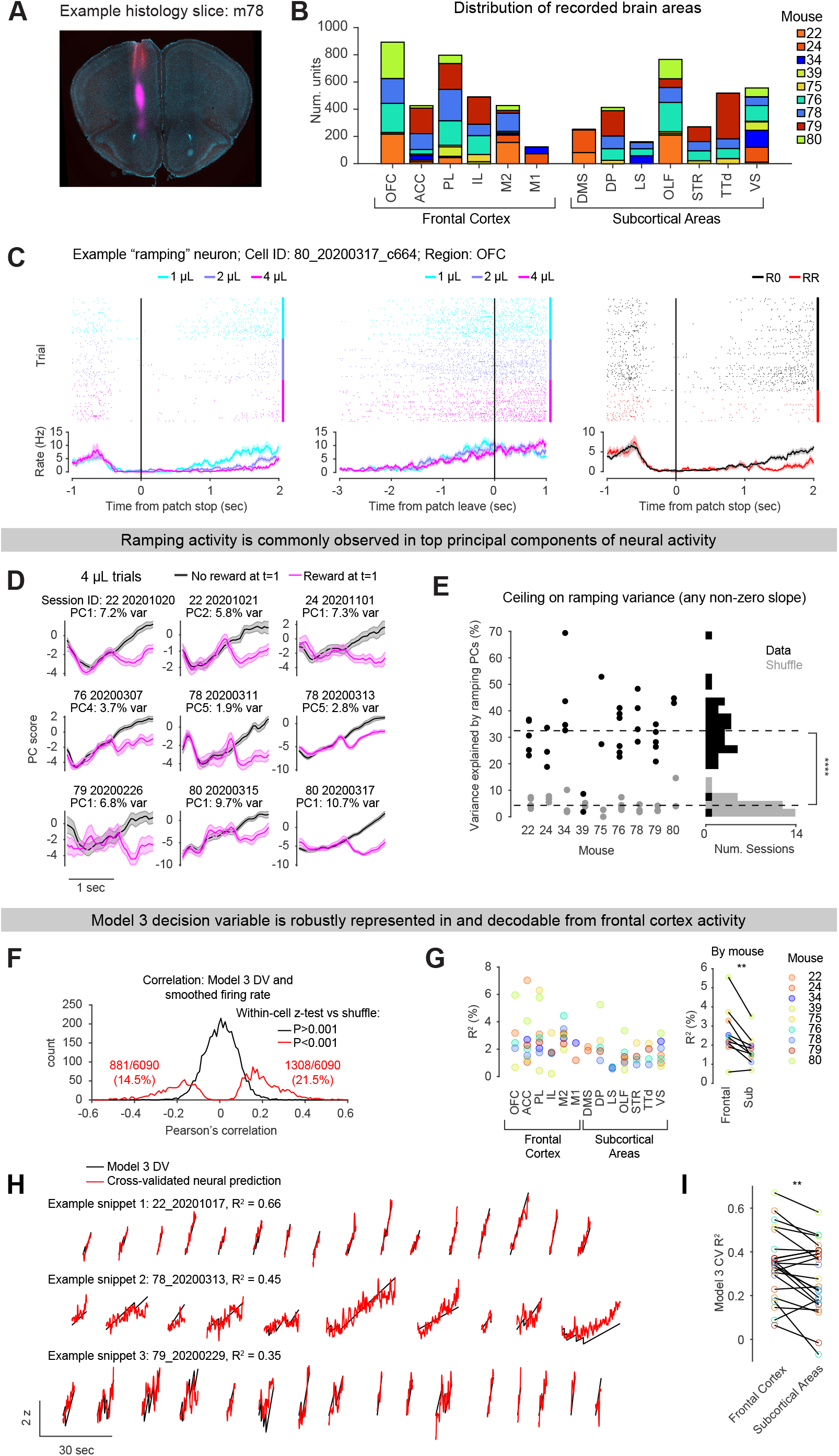
Ramping suppressed by reward is a prominent feature of neural activity and enhanced in frontal cortex. A) Example histology slice showing probe tracks targeting M2 and OFC. Red: DiI, Purple: DiD. Probe tracks were tilted relative to brain slices, so only part of each probe track is visible in each slice. B) Distribution of recorded brain areas from 9 mice. Areas designated as “Frontal Cortex” are grouped together. C) An example neuron showing ramping activity which was suppressed by reward delivery. Left: PSTH aligned to patch stop, split by reward size (cyan: 1 *μL*, purple: 2 *μL*, magenta: 4 *μL*; trials with reward at t=1 omitted). Middle: PSTH aligned to patch leave, split by reward size. Right: PSTH aligned to patch stop, split by whether reward was delivered at 1 second (red) or not (black); rewards of different size combined. D) Hand-picked principal components (PCs) of neural activity showing reward integrator-like activity. For simplicity, only large reward size (4 *μL*) trials are shown. Magenta/black traces are trials with reward/no reward delivery at t=1. Lines indicate means and shaded area indicates standard error of mean. E) PCs with significant ramping slopes, positive or negative, were identified using a shuffle analysis (see Methods and Fig. S6D). The total variance explained by these PCs was computed (black points, each point represents a session). For a shuffle control (gray points), neural activity traces were randomly rotated relative to task events, and the same analysis was performed, once per session. Because PCs with any non-zero ramping slope were included, not necessarily ramping patterns matching those in panel D, this analysis gives a ceiling on ramping variance. **** P < 0.0001, data versus shuffle, sign rank test (N = 33 sessions). F) Histogram of Pearson’s correlation between smoothed firing rate and the Model 3 decision variable (DV; Fig. 3) for each neuron in our data set. A shuffle distribution was generated separately for each neuron and a z-test was used to identify neurons with significant correlations (P < 0.001). G) *R*^2^ values for the regression of Model 3 DV on individual neurons’ firing rates (as in E), by brain region. In the left plot, each point represents a recording session. In the right plot, each point represents a mouse. Consistently across mice, frontal cortex areas had higher *R*^2^ values than sub-cortical areas. ** P < 0.01, paired t-test (N = 9 mice). H) Snippets of several contiguous patches from three example recording sessions, showing the Model 3 DV in black and the cross-validated neural prediction in red. Neural predictions were generated using linear regression on training trials applied to held-out test trials. I) Comparison of Model 3 DV coding between Frontal Cortex and Subcortical Areas. For each within-session comparison, neurons were down-sampled so that each region had the same number of neurons. Only sessions with at least 20 neurons in each of the two brain regions were kept (23/33 sessions). Each pair of data points represents a recording session, and colors represent mice. ** P < 0.01, paired t-test (N = 23 sessions). **Frontal Cortex Areas:** OFC: Orbitofrontal cortex, ACC: Anterior cingulate cortex, PL: Prelimbic cortex, IL: Infralimbic cortex, M2: Secondary motor cortex, M1: Primary motor cortex. **Other Areas:** DMS: Dorsomedial striatum, DP: Dorsal peduncular area, LS: Lateral septum, OLF: Olfactory areas, STR: Striatum, TTd: Taenia tecta dorsal part, VS: Ventral striatum.

Like the example neuron, the top principal components (PCs) of neural activity on patches often resembled an upward ramp with interim drops at reward events (Fig. 4D). We identified “ramping” PCs as those having significantly non-zero slope between 0.5 and 2 seconds on trials with no reward at t=1 (see Methods). The average total variance explained by ramping PCs was 32.5 ± 2.2% (n = 33 sessions), significantly greater than a shuffle control in which neural traces were jointly circularly permuted (4.3 ± 0.5%, paired t-test p < 10e-14; Fig. 4E).

The prevalence of reward-modulated ramping activity suggests that DVs derived from behavioral models may be directly linked to spiking activity. Indeed, the activity of individual neurons was significantly correlated with ramping DVs derived from behavioral models (Model 3 DV; Fig. 4F). Correlations could be either positive or negative but were more frequently positive than negative (1,308 and 881 cells out of 6,090 with z-test, p < 0.001 versus shuffle, respectively; Fig. 4F). Notably, Model 3’s DV explained more variance in single neurons’ firing rates in Frontal Cortex compared to Subcortical Areas, suggesting a specialized role for Frontal Cortex in this task (Fig. 4G).

To quantitatively measure the degree to which DVs were represented in neural population activity, we fit cross-validated linear models predicting each DV from the joint activity of co-recorded neurons (Fig. 4H, Fig. S4B). DVs could be reliably decoded from neural activity in most sessions (Fig. S4C). Although Model 3 best explained mouse behavior (Fig. 3), DVs from Model 2 could be decoded from neural activity with similar fidelity, and Model 1 with only slightly lower fidelity, consistent with a recent report^27^ (mean CV *R*^2^ ± SEM for Model 1 = 0.31 ± 0.03, Model 2 = 0.38 ± 0.04, Model 3 = 0.35 ± 0.03; n = 25 sessions after excluding 5 sessions with CV *R*^2^ < −0.2, see Methods; Fig. S4D). Moreover, Model 3 DV decoding from neural activity was unrelated to performance of the behavioral fit on a per-session basis (Fig. S4E).

To assess how the ability to decode the Model 3 DV relates to brain region, we chose sessions in which we had sufficient neurons in both Frontal Cortex and Subcortical Areas (least 20 neurons each).

Units were down-sampled so that both areas contained the same number of neurons in this analysis. In this within-session, unit-number-matched comparison, we were better able to decode the Model 3 DV from Frontal Cortex than Subcortical Areas (Fig. 4I; mean CV R^2^ ± SEM, Frontal Cortex = 0.34 ± 0.03, Subcortical Areas = 0.28 ± 0.03, paired t-test, p = 0.0037, n = 23 sessions after excluding 7 sessions with insufficient neurons and 3 sessions with CV *R*^2^ < −0.2 for both Frontal Cortex and Subcortical Areas).

Importantly, DVs were also correlated with behaviors such as running speed and lick rate (Fig. S4F). These correlations were generally lower in patches in which mice resided longer but remained non-zero (Fig. S4G). Such correlations may reflect embodied cognition and therefore should not necessarily be viewed as confounds per se (see Discussion). Nevertheless, to rigorously isolate task-related activity from correlated behavioral variables, we next turned to a multivariable regression approach using generalized linear models (GLM).

### GLM modeling with unsupervised clustering reveals six clusters of neurons, the most prominent of which shows ramping activity and reward responses with opposite signs

To comprehensively characterize the space of task-related neural activity, we used a Poisson Generalized Linear Model (Poisson GLM; Fig. 5A, top; Fig. S5A; snippet of GLM fit to an example neuron, Fig. S5B,C) which included the following regressors: session time and its square, to capture slowly varying changes in firing rate; behavioral variables derived from the rotation of the running wheel (position, speed, and acceleration) and the lick sensor (lick rate and its derivative); discrete events at patch stop, patch leave, and reward times, which were convolved with a raised cosine basis, separately for each reward size; and three ramping decision variables per reward size which could be used to construct the decision variables in the behavioral models (Time on Patch, Total Reward, and Time Since Reward). In total, the GLM included 85 variables. Elastic net regularization with 90% L1 (regularization strength selected through cross-validation, see Methods) encouraged the model towards sparsity, thus mitigating issues related to correlated regressors. As a further test of whether behavioral variables such as running speed may contribute to ramping activity in our data, we fit also GLMs to the neural data in inter-trial intervals (ITIs), and found that GLM coefficients for running speed were more correlated between odd and even patches than between odd and even ITIs, suggesting that even if ramping activity is correlated with speed it is not a low-level, non-specific motor code that is common to both patches and ITIs (Fig. S4H).

**Figure 5:**
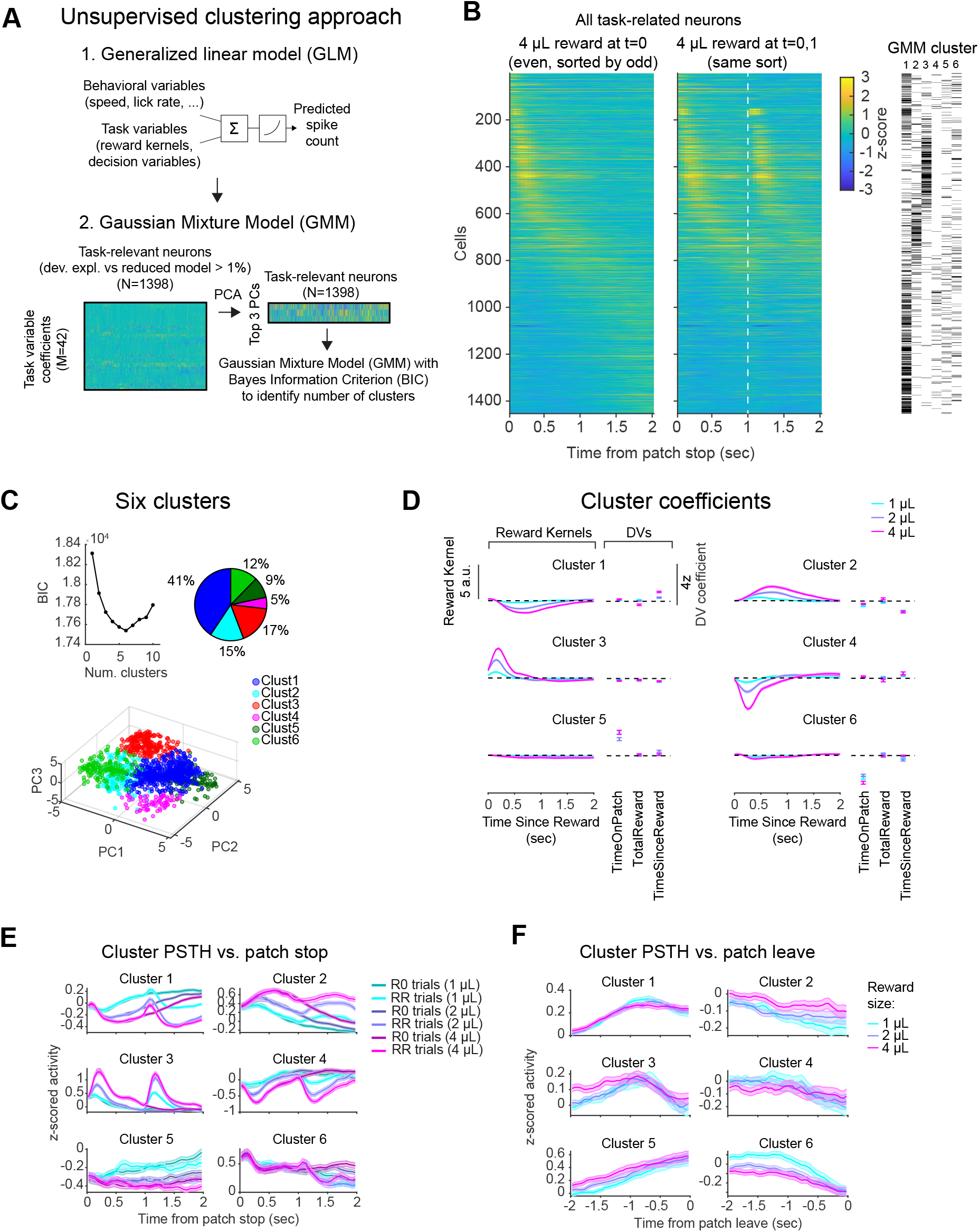
Unsupervised clustering reveals six clusters of neurons, the most prominent of which shows ramping activity and reward responses with opposite signs. A) Schematic of the analysis approach: A Poisson GLM to estimate task variable coefficients, which are used to cluster neurons using a Gaussian Mixture Model (GMM). B) Z-scored neural activity for all task-related neurons from all brain regions on “40” trials (4 *μL* reward at 0 seconds, no reward at 1 second; left panel) or “44” trials (4 *μL* reward at 0 and 1 second; right panel; white dashed line indicates reward at 1 second). Neurons were sorted based on the time of peak activity on odd 40 trials; the left panel showed the same sort on even 40 trials, and the right panel shows the same sort on all 44 trials. An initial transient reward response (as in Fig. S4A) gradually transitioned into upward ramping activity which was suppressed by reward delivery (as in Fig. 4C). GMM cluster identity for each neuron is indicated on the right. C) GMM clustering on the top 3 PCs of task variable coefficients was used to identify clusters of neural activity patterns. **Top left:** Bayesian information criterion (BIC) was used to select the number of clusters (minimum BIC: 6 clusters). **Top right:** Percentage of neurons assigned to each cluster. Clusters were ordered so that patterns with similar shapes but opposite signs were adjacent (see panel E). **Bottom:** Task-related neurons projected into the PC space used for clustering, colored by assigned cluster. D) Average z-scored GLM task variable coefficients for each cluster. Reward kernel coefficients were multiplied by corresponding basis functions and summed to generate the predicted reward response. E) Average PSTHs of z-scored neural activity following patch stop for each cluster, split by reward size and whether reward was delivered at 1 second (R0 indicates a reward at 0 second, not 1 second, RR indicates rewards at both 0 and 1 second). F) Average PSTHs of z-scored neural activity aligned to patch leave for each cluster, split by reward size.

To assess model performance, we computed cross-validated percent deviance explained compared to a null model (Fig. S5D, top; mean % deviance explained ± SEM = 5.2 ± 0.1, n = 6,090 neurons). Reward kernels and decision variables were termed “Task Variables” and used to identify task-related neurons. Percent deviance explained was computed for full models compared to models without Task Variables (“reduced models”), and neurons with deviance explained > 1% in this comparison were deemed “task-related” (Fig. S5D, bottom; 1,458/6,090 neurons, mean % deviance explained versus reduced model ± SEM = 0.87 ± 0.03). Mirroring Model 3’s DV decoding performance, GLM performance was higher for Frontal Cortex areas than other areas (Fig. S5E, mean % deviance explained ± SEM, Frontal Cortex = 6.5 ± 1.0, Subcortical Areas = 4.1 ± 0.6, paired t-test, p = 0.016, n = 9 mice). Aligning the activity of task-related neurons to patch stop, splitting trials by those in which a reward was delivered at 1 second or not (RR versus R0 trials), and sorting by the time of peak response revealed a “wave” of activity following each reward: A rapid reward response with variable latency which gradually transitioned into upward ramping activity that was reset by reward delivery (Fig. 5B).

We clustered task-related neurons based on their GLM coefficients to characterize the space of activity patterns in our task in an unbiased manner (Fig. 5A, bottom). Clustering was performed on a dimensionality-reduced matrix of task-variable coefficients by neurons (top 3 principal components, 34.7% of variance; Fig. S5F), with number of clusters (K = 6) chosen to minimize the BIC (Fig. 5C) (Methods). The resulting 6 clusters could be grouped into 3 pairs with similar shapes but opposite signs (Fig. 5D): (1) A pair of clusters with a slow negative/positive reward response, scaled by reward size, and positive/negative coefficients on Time Since Reward (Clusters 1 and 2), (2) A pair of clusters with rapid reward response, scaled by reward size, and minimal ramping coefficients (Clusters 3 and 4), and (3) a pair of clusters with minimal reward coefficients but significant positive/negative coefficients on Time On Patch (Clusters 5 and 6). Similar clusters could be identified in 6 out of 9 mice; the other 3 mice had too few task-related neurons to reliably identify the same patterns (Fig. S5G). PSTHs of the average activity of each cluster aligned to patch stop and split by reward delivery at one second matched the patterns of GLM coefficients (Fig. 5E). Moreover, aligning to patch leave revealed that Cluster 1 ramped to a common threshold approximately 0.5-1 seconds prior to patch leave (Fig. 5F). Because of the clear ramp-to-threshold-like properties of Cluster 1, we considered this to be our “ramping” population (568/1,398 task-related neurons).

A greater percentage of neurons in frontal cortex areas was identified as “task-related” compared to other areas (Fig. S5H; frontal cortex areas: 916/3,156 [29%], other areas: 482/2,934 [16%]), consistent with a specialized role for frontal cortex in our task. However, of task-related neurons, similar fractions were assigned to Cluster 1 across brain areas (Frontal Cortex: 40.7%, Subcortical Areas: 40.5%). To assess whether our GMM Clusters may be enriched for certain cell types^29,30^, we classified waveforms as Regular or Narrow based on spike width (Fig. S5I). In Frontal Cortex recordings, GMM Cluster 3 (transient positive response to reward) contained an over-representation of Narrow waveforms, indicating that our functionally defined clusters may map onto cell type-specific populations of neurons in the cortex.

### The ramping population exhibits ramp-to-threshold activity with reward sensitivity modulated by latent state

We next examined the activity of our “ramping” population (Cluster 1) more closely. Splitting trials by PRT revealed that the activity gradually ramped to a common threshold across trials of differing duration (Fig. 6A, Fig. S6A,B). While the magnitude of the reward-induced dip was strongly modulated by reward size (Fig. 6B), the ramp slope could be modulated positively or negatively across individual neurons, resulting in an average modulation not significantly different from zero (Fig. 6C). We therefore looked specifically for task-related neurons with ramping slopes modulated by reward size and identified a subgroup of neurons with this property (362/1398 neurons, 26%; Fig. S6F-I). Of these, 156 neurons (43%) showed modulation opposite to the direction of the ramping slope itself (i.e. shallower slopes for larger reward size), as in the behavioral model (Fig. S6F). Notably, though trials with reward at t=1 were not used to identify these neurons, their activity dipped following reward delivery at t=1, like the behavioral DVs. Applying a similar analysis to neural PCs, we found that ramping PCs with ramp slope modulated by reward size accounted for ∼10% of in-patch variance (Fig. S6J), roughly 1/3 of the total variance explained by ramping PCs (Fig. 4E). These activity patterns provide a plausible neural basis for an integrate-to-threshold process guiding patch foraging decisions.

**Figure 6:**
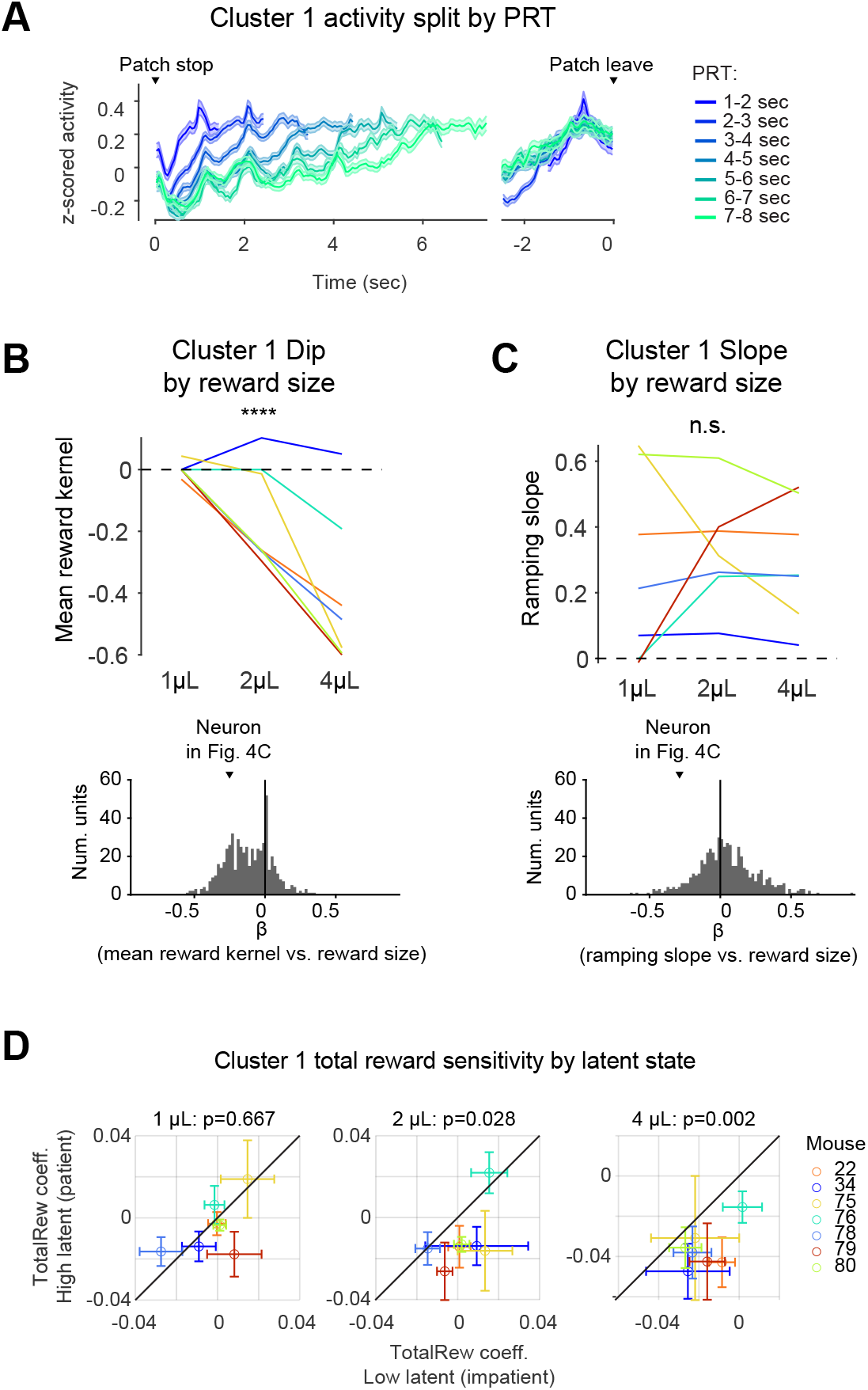
The ramping population (Cluster 1) exhibits ramp-to-threshold activity with reward sensitivity modulated by latent state. A) Average PSTHs of z-scored neural activity for each cluster, split by patch residence time. Lines indicated means, shaded regions indicate standard error of the mean (N = 568 Cluster 1 neurons). B) **Top:** Mean reward kernel GLM coefficient for each reward size, by mouse. ****, P < 0.0001, fixed effect of reward size, linear mixed effects model with a random effect per mouse (N = 7 mice). **Bottom:** Histogram of regression coefficients (*β*) of mean reward kernel versus reward size, for all Cluster 1 neurons. The example neuron shown in Fig. 4C is indicated. In the mouse-level analyses in panels B-D, two mice with very few task-related neurons were excluded (mice 24 and 39, with 30 and 17 task-related neurons respectively), leaving 7 mice. C) Same as B, but for the slope of ramping activity, extracted from trials with no reward at t=1 (see Fig. S6D). n.s., not significant, fixed effect of reward size, linear mixed effects model with a random effect per mouse (N = 7 mice). D) Total reward (TotalRew) GLM coefficients for Cluster 1 neurons, from GLMs fit to high and low latent trials separately. TotalRew tracks the total number of rewards received on a given patch (see Fig. S5B for example traces). P-values are from paired t-tests on the average coefficient in high and low latent trials by mouse (N = 7 mice; not corrected for multiple comparisons).

Because latent state (or “patience”) strongly influenced mouse behavior (Fig. 2), a natural question is whether latent state affects the coding properties of ramping neurons in a manner that could explain the behavioral changes. To test this, we fit neural GLMs to high and low latent trials separately (trials split in half by latent state within each recording session), following the same GLM fitting procedure as above. Cluster 1 neurons’ *TotalRew* coefficients were more negative in high latent (patient) trials compared to low latent (impatient) trials (Fig. 6D), consistent with the putative integration mechanism implementing leave decisions. No other decision variable or behavioral variable coefficient, including running speed and lick rate, differed between high and low latent trials (all p > 0.05), but average Cluster 1 reward kernel coefficients were slightly more negative in high compared to low latent trials, consistent with the enhanced sensitivity to *TotalRew* (paired t-tests, 1 *μL*, p = 0.093; 2 *μL*, p = 0.034; 4 *μL*, p = 0.039). Thus, latent state may shape decision making in part by enhancing the sensitivity of cortical ramping dynamics to integrated reward.

### State space modeling demonstrates that neural population activity is ramping, not stepping, on single trials

In previous sections, we identified a cluster of neurons that showed ramping activity on average (“Cluster 1”, Fig. 5-6). We next tested whether single trial dynamics within this sub-population (e.g. Fig. 7A) were also continuously varying ramps or were discrete processes – an issue that has been controversial in decision-related activity in perceptual decisions^31,32^. Although previous analyses relied on single neuron recordings, we reasoned that our ability to record the activity of multiple “ramping” neurons simultaneously may help us resolve this issue for patch foraging decisions^33,34^. To this end, we fit a continuous ramping model and a discrete state stepping model to the neural population response (Cluster 1 spike rate, summed across neurons; Fig. 7B,C). The ramping model had a pulse response to rewards and otherwise ramped in the absence of rewards. The stepping model had two discrete firing rate states; the firing rate started in the low state and probabilistically jumped to the higher state following Markovian dynamics. Additionally, to capture reward responses, the firing rate returned to the low state after rewards, whereas otherwise the firing rate was fixed to remain in the high state. We found that all 16 sessions with at least 10 Cluster 1 neurons were better fit by the ramping than the stepping model (Fig. 7D). Additionally, the latent accumulator drop was greater for larger reward sizes (Fig. 7E, left; regression slope of model parameter vs. reward size, mean ± SEM = -0.012 ± 0.0041, t-test, p = 0.0093, n = 16 sessions), and slopes were marginally shallower (Fig. 7D, right; regression slope of model parameter vs. reward size, mean ± SEM = -0.025 ± 0.012, t-test, p = 0.065, n = 16 sessions), consistent with the hypothesized integration mechanism which accounts for the impact of rewards on mouse wait times (Fig. 3).

**Figure 7:**
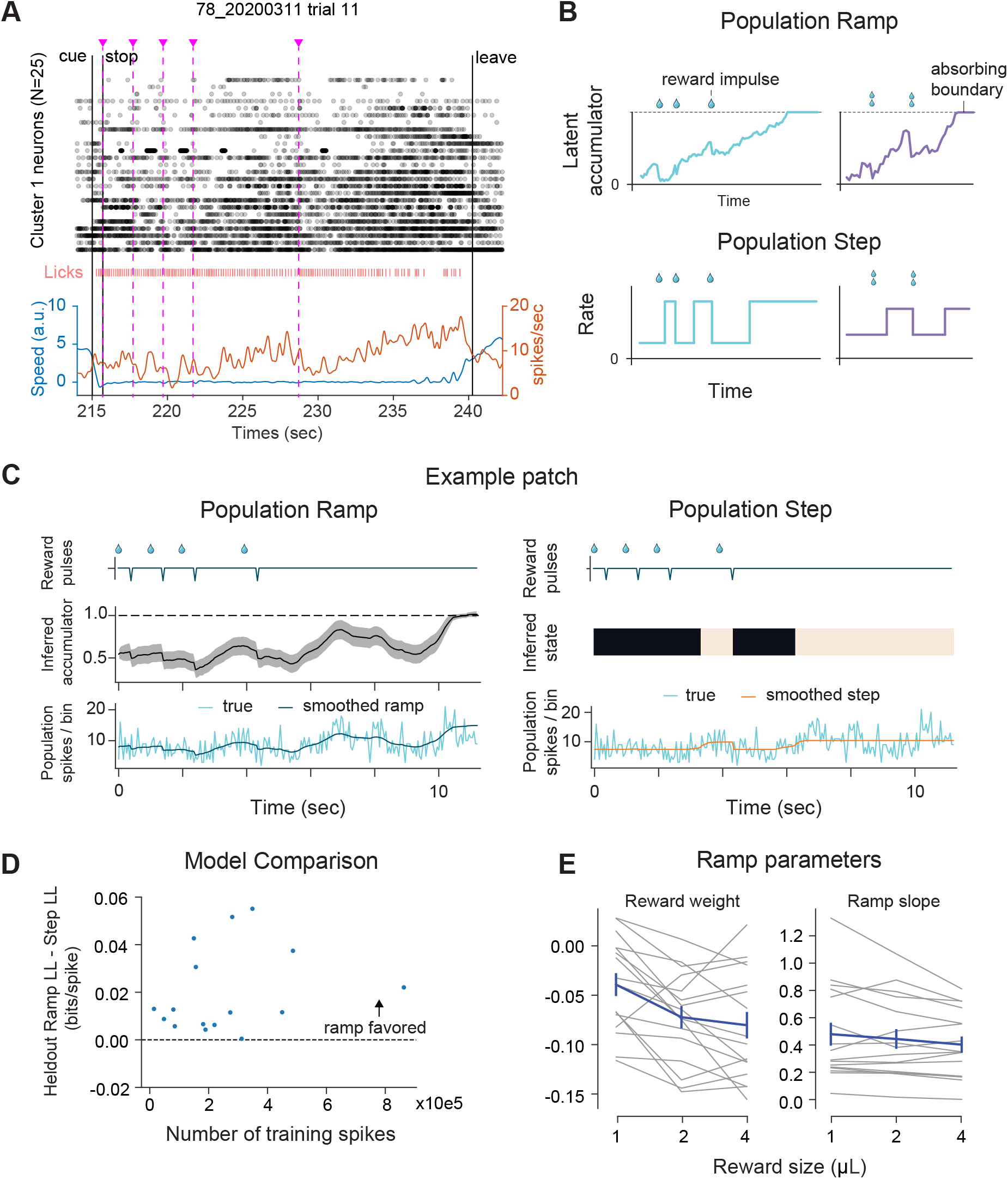
Cluster 1 population activity is ramping, not stepping, on single trials. A) Example trial showing ramping activity of simultaneously recorded Cluster 1 neurons while the mouse remained still on the patch. Top: Raster plot of 25 Cluster 1 neurons. Middle: Raster plot of mouse licks. Bottom: Mouse speed (blue) and average firing rate of Cluster 1 neurons (red). Magenta dashed lines: Reward delivery (4 *μL*). B) Schematic of ramp and step models. Models were fit to Cluster 1 neurons. C) Single trial ramping and stepping model fits for an example session (80_20200317). D) Model comparison across sessions. Only sessions with at least 10 Cluster 1 neurons were included (16/33 sessions). For 16/16 sessions, the ramp model outperformed the step model on held out data. E) Ramping reward coefficients and slopes across reward size, by session. Note that reward weights become more negative with reward size (i.e., bigger dip), resulting in a larger decrease in the tendency to leave.

## Discussion

In the present study, we sought to identify the algorithm underlying leaving decisions in a classic patch foraging paradigm long studied in the framework of optimal foraging theory in behavioral ecology. We modeled an environment with multiple “patches” with depleting resources, using a novel virtual reality-based patch foraging task, which allowed us to precisely control the timing and the amount of reward. The naturalistic decision-making paradigm balanced behavioral complexity with analytical tractability, maintaining a link to ethologically relevant behavior while allowing us to carefully probe its neural and algorithmic basis. We varied patch richness along two independent dimensions (reward size and frequency) and showed that mice’s PRTs were sensitive to both, consistent with a basic prediction of the MVT. However, more direct tests of the MVT, facilitated by the reward schedules and stochasticity in our task, revealed that mouse behavior systematically deviated from MVT predictions: The instantaneous reward rate at leave time differed between patch types, and differences in PRT between patch types increased with mean PRT. While we cannot rule out alternative formulations of MVT that could capture these effects, we found that both are explained by integrator models. Of three integrator model variants, the Reward Integrator (Model 3) best fitted our behavioral data, as confirmed by both quantitative model comparison and diagnostic trial type comparisons. Notably, a model fit with a single set of free parameters shared across the population was able to account for mean PRT variability within and across mice, demonstrating that a common algorithm underlying the choice process could produce the observed wide range of behaviors. This raises the intriguing possibility that scaling of a similar common algorithm could be responsible for variation in human behaviors such as impulse control, and that maladaptive scaling could underlie pathological conditions.

In large-scale electrophysiological recordings across frontal cortex and underlying subcortical areas, we found a prevalence of single neurons whose activity exhibited a ramping profile, which integrated the elapsed time and intermittent rewards—two categorically distinct variables—in an opposing manner, consistent with the Reward Integrator model. We identified a population of “ramping” neurons (Cluster 1) and found that the single-trial activity of these neurons was better described by continuous ramping rather than a discrete stepping process. The reward amounts affected the magnitude of the reward-induced dip, and in some neurons modulated the ramp slope, resulting in population activity that could be used to decode the Model 3 ramping decision variable, more robustly in frontal cortex compared to subcortical areas. Ramping activity persisted for up to tens of seconds, thus demonstrating that the same mechanism could be scaled to produce both patient and impulsive choices. Together, these results indicate that a continuous integration process reflected in the activity of frontal cortex neurons is a likely mechanism by which mice make patch leaving decisions during foraging.

### Integrator models can account for deviations from the Marginal Value Theorem

The MVT provides a normative solution to the question of when an animal should leave a patch of depleting resources. However, such a precise solution is practicable only under idealized conditions^17^ and quantitative tests of the theory reveal a range of contradictions^5,35-39^. While modeling efforts have reconciled such results with MVT by invoking non-linear marginal utility curves^17,38^ or quasi-hyperbolic temporal discount functions^39^, we show that integrators—while operating at the level of algorithm rather than objective—can readily explain deviations from the predictions of MVT in its simplest form.

Moreover, in situations where an MVT theoretic solution is intractable, integrators can provide a means for allocating greater foraging time to locations in which resources are more abundant.

We structured the task such that MVT makes explicit predictions about the differences in relative leave times across patch types. By utilizing a reward schedule with stochastic rewards of diminishing probability, rather than a deterministic reward schedule or one with fully observable decays in reward size, mice were required to make their leaving decisions in the face of uncertainty, and were thus prevented from using a strategy such as leaving once an observed reward dropped below a particular size threshold. By tuning the statistics of the task in a way that allows for tests of explicit predictions from the MVT about differences in PRT across patch types, we were able to observe discrepancies from theoretical expectations that are not readily explained by previous accounts.

For example, **Prediction 3** dictates that, due to the steady decay rate we used for reward probability within patches, mice should remain on patches with double the reward size for 5.6 seconds longer. We observed that this relationship was consistently broken across animals, with mice producing smaller than predicted average PRT differences across reward sizes (Fig. 3C). Temporal discounting would not account for this deviation since discounted reward expectation would scale proportionally across patches of different reward size. A nonlinear expected utility curve across reward sizes could, in principle, account for these deviations by scaling the marginal gains in expected value from larger rewards sublinearly. However, we also observed that the magnitude of deviations in expected reward rate at the time of patch leave across reward sizes was not constant across time and subjects. Instead, PRT differences across reward sizes were dependent on latent levels of patience both within and across subjects (Fig. S2A,E), thus violating **Prediction 2**. Critically, this would require the shape of the utility curve across reward sizes to *depend on patience* to explain the observed PRTs. An account of this dependency, or analogous dependencies for alternative explanations such as risk sensitivity or reward size uncertainty, would therefore be needed. In contrast, these deviations can emerge naturally from integrator processes.

### Latent scaling of a common algorithm explains PRT variability across mice

Decision making varies dramatically with the internal state of the organism^40-44^. While mice were value-based in their PRT behavior, they exhibited substantial variability in their overall willingness to wait (“patience”). Interestingly, we observed that reward sensitivity scaled systematically with patience, resulting in greater differences in PRT as a function of patch value when patience was higher. Coefficients for reward size and frequency effects on PRT varied widely, but they did so with approximately the same ratio across mice (Fig. S2A). Because of this common scaling, we were able to consistently capture PRTs across conditions and individuals with a GLME where reward effects were fixed across the population, and with only a random effect term per mouse to scale as a multiplier (Fig. 1G). Furthermore, replacing the per-individual random effect with a term for latent patience scaling allowed us to robustly predict PRTs with all model parameters shared across mice (Fig. 2D). These results suggest that mice with different PRT behavior may employ a common algorithm, with individual differences in scaling.

Consistent with this hypothesis, we were able to robustly predict single trial PRTs across the population with our competitive integrator model using a shared set of parameters, with the only source of individual differences arising from the scaling introduced by the patience estimates (Fig. 3F-G).

While the current experiments do not provide direct insight into what factors contribute to the varying degrees of patience, we can rule out several simple explanations based on the observed mice behavior. Satiation would be a sensible candidate hypothesis, since changes in this state occur over the course of a session, as does the varying latent state. However, the within-session dynamics we observed were complex and heterogenous across sessions, and even the direction of the change varied (Fig. 2C). For the same reasons, we can rule out other variables that vary monotonically, such as physical exhaustion due to running on the treadmill. Moreover, the greatest variance we observed in latent patience estimates were across sessions and mice, whereas we would expect the greatest differences in satiation to occur between the beginning and end of each session since supplemental water amounts were calibrated to keep mouse weights steady throughout training. Therefore, although satiation inevitably plays an important role in varying motivational state, it is also clearly not the sole factor, and its effect likely interacts with other factors resulting in different net effects on behavior across mice or at different times.

The latent state we call patience is related to a broader literature on task engagement, arousal, and motivational state^45-51^ and the idea that decision processes may be non-stationary^44^. Whereas other studies have focused on identifying discrete or abruptly changing latent behavioral states^45,51-53^, ours is a continuous, slowly changing variable^50^ derived from the average of surrounding trials. The patience state is likely influenced by neuromodulation, for example from serotonin^54^ or noradrenaline^43^. In a recent study, optogenetic activation of serotonin neurons increased “active persistence” during foraging^54^, whereas here patience reflects a passive process of waiting for rewards^55^, making it difficult to generalize across studies. As our work focused on spiking activity in frontal areas, further experiments are needed to identify the contributions of neuromodulation to the latent patience state identified here.

In our behavioral modeling, we were best able to capture the dependence of PRT on latent state by using latent state to scale ramping slope (Fig. S3A). In contrast, the primary effect of latent state observed in our neural data was a scaling of the reward-triggered decrements of the ramps (Fig. 6).

Importantly, however, we observed the greatest variance in latent estimates across sessions and mice (Fig. S2C), but could only examine the relationship between neural activity and patience within individual sessions. Therefore, future work tracking the same neurons across sessions would be needed to better understand how patience modulates ramping neural activity.

### Comparison with reinforcement learning models of foraging behaviors

Previous works have analyzed foraging behavior in the framework of reinforcement learning (RL)^38,56-58^. While we did not implement RL models here, integrator and RL models share key commonalities. For one, variables in RL models often ramp over time; for example, the value of staying on a patch decays over time and increases when rewards are delivered, mirroring our ramping DVs. It therefore may be difficult to distinguish RL models from integrator models using ramping neural signals alone. Indeed, integrator models may be seen as a kind of heuristic implementing an RL model, with the value of the integrator representing the relative value of staying on the patch. In general, integrator models are a highly flexible model class capable of capturing a range of behaviors, and previous models including MVT and RL variants may be mapped to integrators by specifying the rules which set and update integrator parameters^17^.

As in Shuvaev et al.^56^, we found that animals left richer patches at higher reward rates, which is suboptimal compared to ideal MVT (Fig. 3A). Shuvaev et al. account for this using R-Learning, an RL algorithm which updates patch value and environment value in parallel and compares them to determine the probability of patch leave. In this framework, mice leave good patches too early because the estimate of environment value has also increased, making leaving more attractive. In contrast, we account for this deviation from optimality empirically by fitting different slope parameters for each reward size (Fig. 3E).

Interestingly, our models do not explicitly predict overstaying on patches with low reward sizes and understaying when rewards are larger. Instead, they dictate that this trend is contingent upon the latent patient state because reward size-dependent differences in slope accumulate over time (Fig. 3K). At levels of patience higher than those generally observed in our experiments, the models predict this trend would reverse, with subjects instead waiting too long on patches with large rewards relative to those with smaller rewards. Future experiments could test this prediction.

Notably, over up to 30 days of training, we observed no change in relative PRT across patch types, nor did we observe an increase in average reward rate (Fig. S1F). This suggests that animals are not gradually learning the task’s reward statistics, potentially ruling out RL models that feature such slow learning. Rather, the data are most consistent with mice implementing a default integration strategy which may be adjusted within sessions by local reward history and other factors, such as patience. Indeed, we found a small tendency for mice to wait longer on patches when previous patches were smaller reward size (Fig. S1G). This may indicate that a local integration of reward history sets a threshold for patch leaving which tracks environment value^38,56^. Future work can investigate whether and how trial history modulates the parameters of the integration models proposed here.

### Implications of correlated behavioral and cognitive variables

Cognitive variables are notoriously difficult to separate from behavior in animal experiments. As shown in Figure S4, several behavioral variables (most prominently position, speed, lick rate) correlated either positively or negatively with ramping decision variables, raising the possibility that some of the ramping activity in our neural data could be associated with task-related movements. Indeed, several recent studies have reported movement-related activity widely throughout the brain, beyond classically movement-related areas^59,60^. Such correlations likely contribute to the ramping signals in our data, especially for mice in which these correlations were high. However, movement is unlikely to fully account for ramping activity, for the following reasons: (1) Ramps were observed even in trials in which mice remained completely still on the running wheel (Fig. 7A), (2) A population of ramping neurons (“Cluster 1”) could be identified from GLM regression coefficients, in which movement-related activity had been regressed out by including behavioral variables in the model (Fig. 5, 6), (3) Cluster 1 firing rates preceding leave time were similar across reward sizes despite different leaving speeds (Fig. 5F, Fig. S1B), and (4) Speed coding was qualitatively distinct in the trial periods and inter-trial-intervals (Fig. S4H). An additional possibility is that correlations between decision variables and behavior could, in part, reflect an embodied cognitive process^61^. The extent to which behavior is itself an element of natural decision-making represents an interesting direction for future study.

Although Model 3 best explained mouse behavior in our task, decision variables from all three models could be decoded from neural activity with similar accuracy (Fig. S4D). This could reflect a “reservoir” of decision variables in frontal cortex, in which strategies that are not currently used are nevertheless continuously tracked in neural dynamics^27^. However, caution must be taken to consider the contribution of task-related movements to these results. For example, licking patterns are similar following each reward and may contribute more to a “Model 2” (reset)-like pattern of neural activity, regardless of the actual decision algorithm implemented by the brain. If we simply exclude *all* movement-related activity (which is not practical), then we risk discarding activity related to embodied cognitive processes. At the same time, it is circular to identify movements as “embodied cognition” because they match a particular model, and then exclude other movement-related activity and draw conclusions based on the match between activity and behavior. Thus, we believe the neural data are of limited use in identifying the exact strategy employed by animals in this task. Instead, our argument for Model 3 is behavioral (Fig. 3), and we show that activity related to the decision variable from this model is *present* in neural activity (Fig. 4F-I), providing a potential substrate for patch foraging calculations in the brain.

### Ramping activity during decision making

Ramping activity has been shown to be a prevalent feature of perceptual decision making across a range of organisms and brain areas^13,34,62-64^. It provides a mechanism for accumulating evidence over time^13,34,65^, making relative value decisions^66^, tracking time for episodic memory^67^, and determining action timing^68-72^. We observed similar ramping dynamics within the context of naturalistic foraging, which can itself be conceptualized as a type of evidence accumulation problem^17^, with time and rewards providing evidence for and against leaving, respectively. The flexibility of ramps to scale their slope and integrate categorically distinct inputs, taken in the broader context, suggests that ramping activity could be a ubiquitous feature of decision processes that unfold gradually over time.

## Supplementary Figures

**Figure S1:**
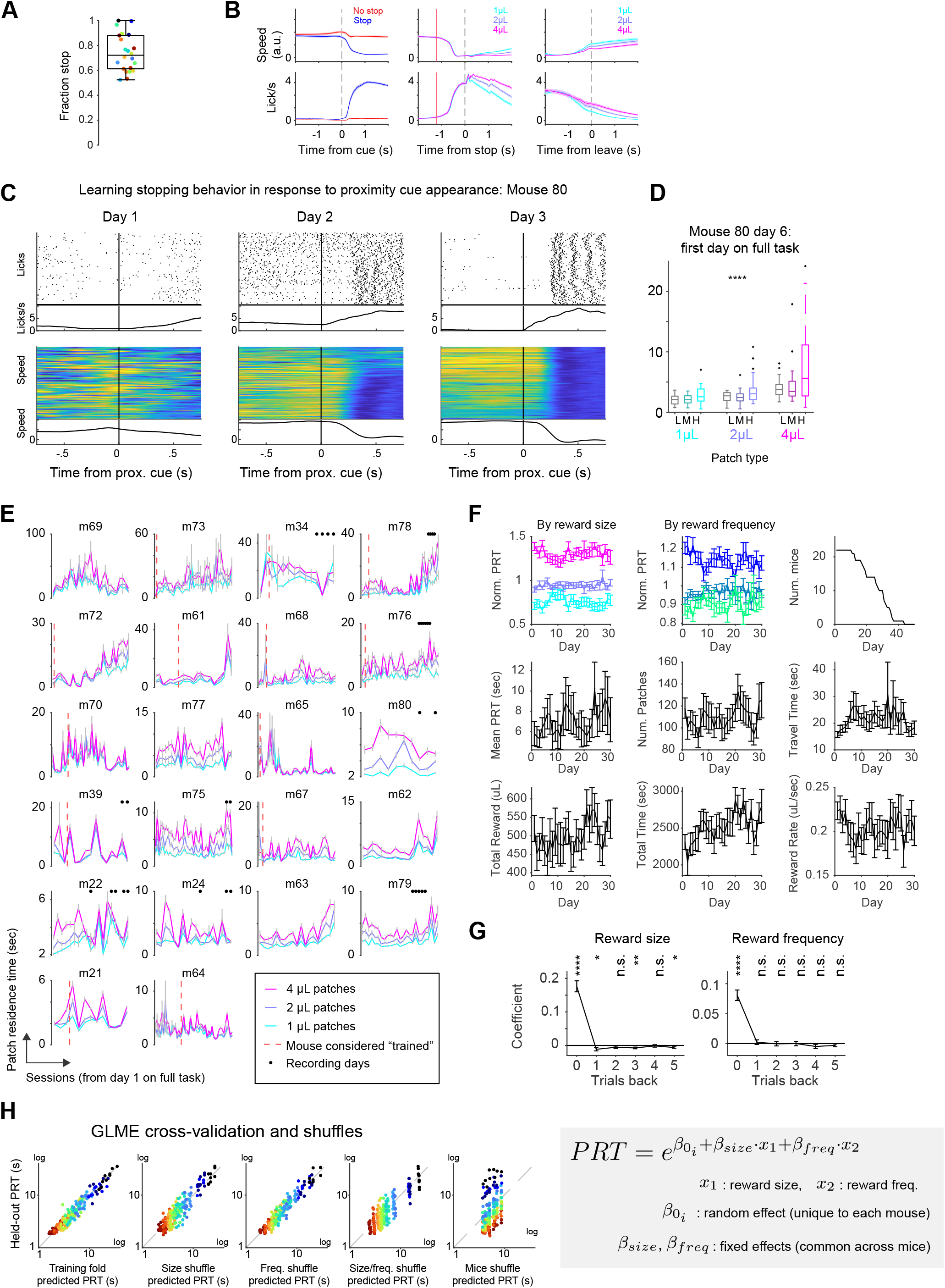
Mouse behavior over training, trial history effects, and task engagement (related to Figure 1). A) Mice stopping behavior in response to the proximity cue. B) Mean population running speed (Top) and lick rate (Bottom) in response to task events. (Left) Aligned to proximity cues appearance, split by trials in which mice successfully stopped (blue) or skipped (red) the patch. (Middle) Aligned to patch stop, split by reward size. Red line indicates mean time of proximity cue appearance. (Right) Aligned to time mice left the patch, split by reward size. C) Example mouse learning over first three days of pre-training in which proximity cue onset is followed by automatic reward delivery after a short delay. Patches are not introduced yet. Mouse learns that proximity cue is predictive of reward and begins showing anticipatory licking and slowdown in response to it. D) First session full task with patches introduced, example mouse. Mice must stop in response to the proximity cue to enter a patch and earn rewards (p < 0.0001, linear regression of PRT on patch value). Column groupings separate patches by reward size. Within columns, patches are split by reward frequency. E) Per subject PRTs across sessions, split by patch reward size. Error bars indicate standard error of the mean. Dashed red line indicates when mice task performance achieved inclusion criteria, after which sessions are included in the final data set. Mice with no dashed red line met inclusion criteria on the first day of the full task. Black dots mark recording sessions included for neural analysis. F) Behavioral performance metrics over training. Norm. PRT = normalized PRT (PRT divided by mean PRT within session). Error bars indicate standard error of the mean over mice. Mice were trained for differing numbers of days; top rightmost plot shows the number of mice remaining on each training day. G) Linear regression of normalized PRT (PRT divided by mean PRT, within session) on reward size or frequency on the current and previous trials. Reward size was 1, 2, or 4 *μL*, reward frequency was coded as 1, 2, or 4 (low, medium, or high), and intercept terms were included in the model. Models were fit using the function fitlm in MATLAB, separately for each session, then coefficients were averaged by mouse. Lines and error bars show mean regression coefficients and standard error of the mean over mice (n = 22). n.s. Not Significant, * p < 0.05, ** p < 0.01, **** p < 0.0001, one-sample t-tests of model coefficients versus zero (n = 22 mice for each test, each lag tested separately, no correction for multiple comparisons). H) Cross-validation of the GLME from Fig. 1G showing held-out PRT versus training fold predicted PRT. Training fold in the leftmost panel is unshuffled. Training fold trial labels for the remaining four were shuffled by reward size, frequency, size and frequency, and mouse ID, from left to right, respectively. Folds yielding the median mean squared error between predicted and held-out PRTs are shown for each shuffle type.

**Figure S2:**
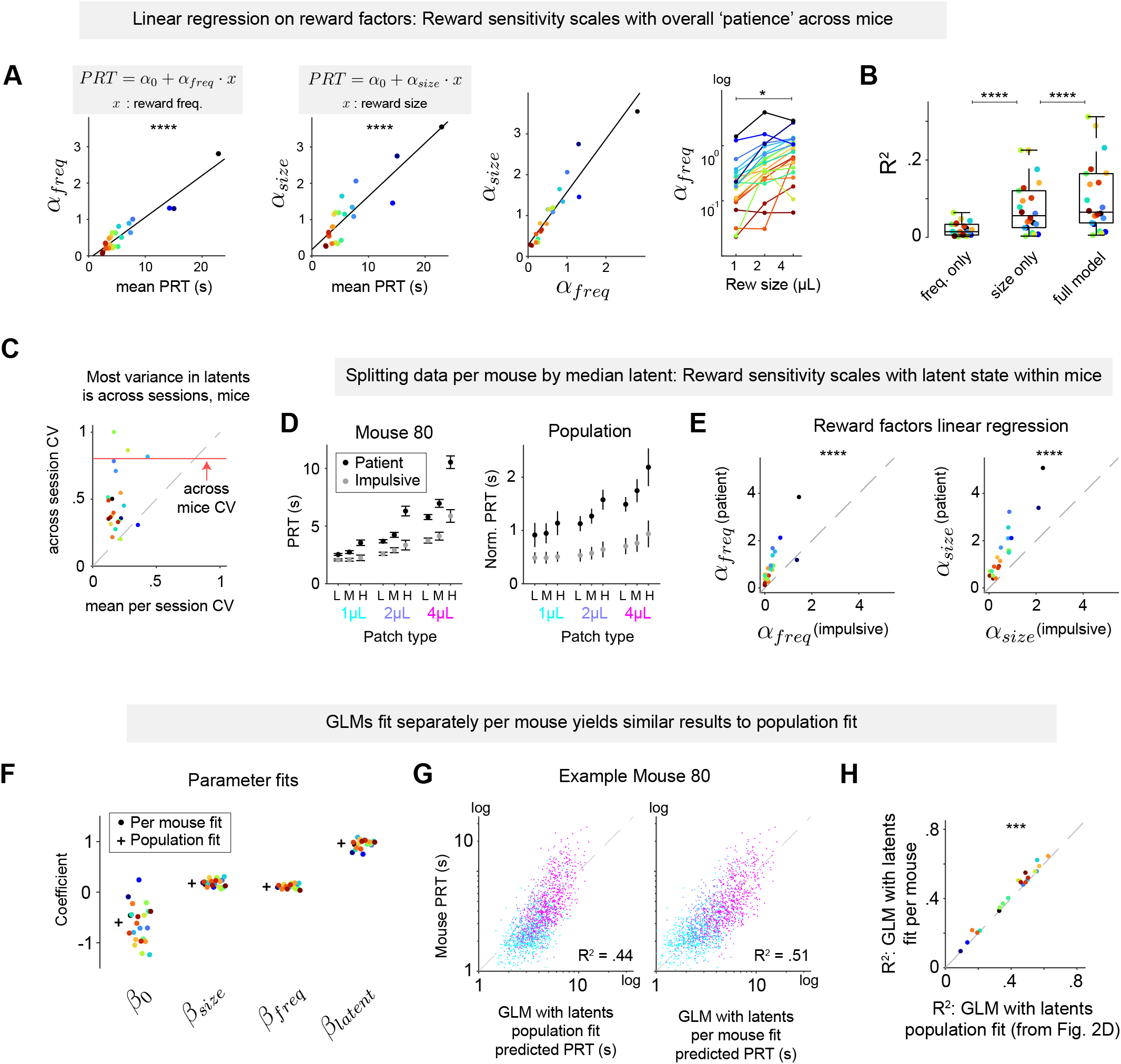
Reward sensitivity scales with patience across and within mice (related to Figure 2). A) Linear regressions of PRT on reward factors per mouse. Left: Regression coefficients for reward frequency and size scales with mean PRT. Middle: Effects of reward size and frequency scale with same ratio across mice. Right: Regression coefficients for PRT on reward frequency are larger for larger reward sizes. B) *R*^2^ values for linear regression across mice for models including reward frequency, reward size, and a full model including both frequency and size as well as their interaction. Stars indicate significance of pairwise Wilcoxon signed-rank tests (both p < 0.0001, Bonferroni-adjusted). C) Comparing coefficient of variance on latent patient estimates within versus across sessions. Red line indicates the coefficient of variance over mean patience estimates across mice. D) PRT per patch type with trials split into two latent state groupings: impulsive and patient, separated by median latent state, example mouse (Left) and population means (Right). E) Linear regressions on reward factors, fit separately per latent state grouping across subjects (p < 0.0001, Wilcoxon signed-rank tests). F) GLM with scaling by latent patience estimates, same as Fig. 2D except fitting with a separate set of parameters per mouse. G) Single trial PRT predictions from GLM population fits (Left) and per mouse fits (Right), example mouse. H) R^2^ statistics for GLM fits per mouse yield modest improvements over population fit (p = 0.0004, Wilcoxon signed-rank test, Bonferroni-adjust).

**Figure S3:**
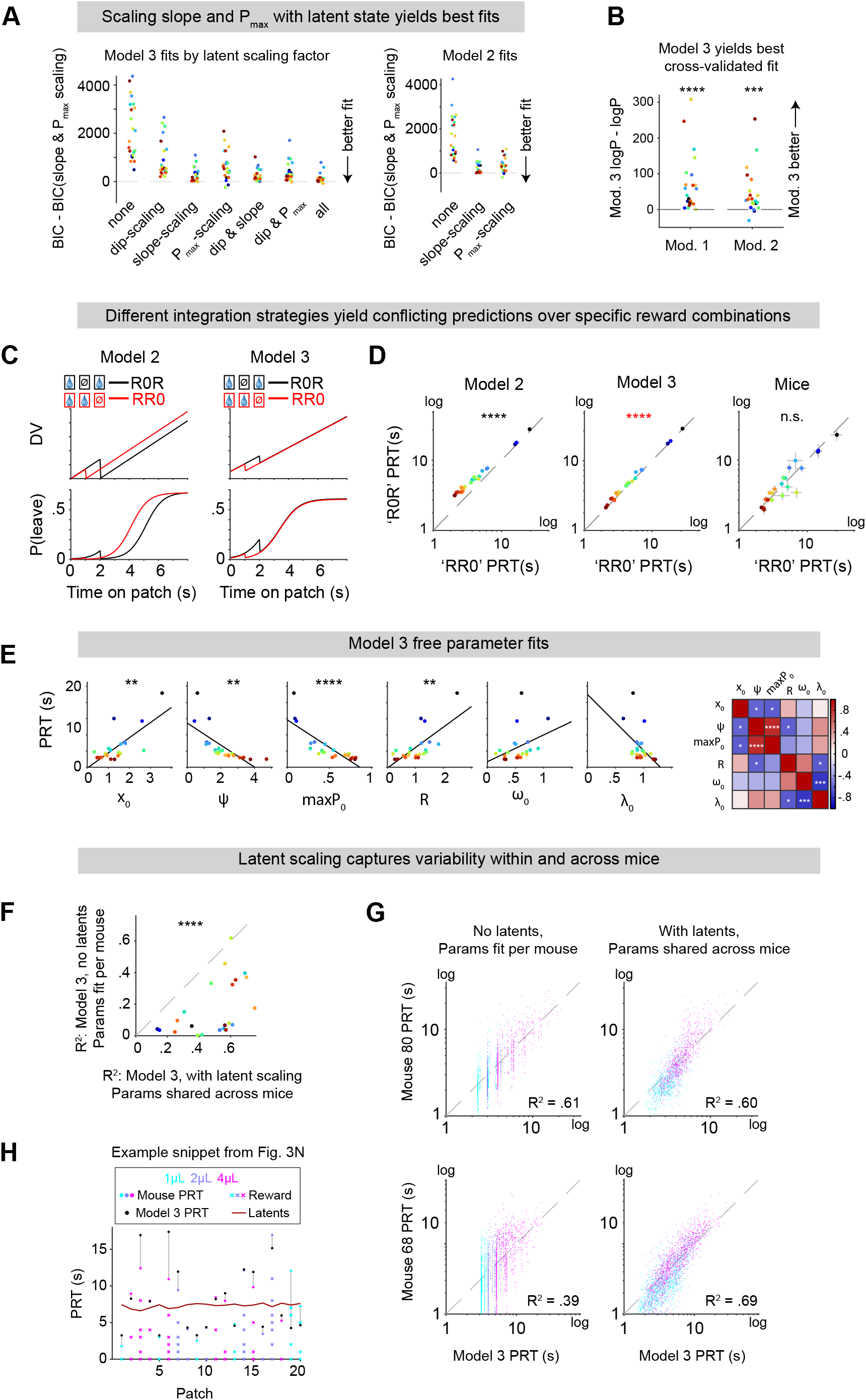
Additional analyses of behavioral model fitting (related to Figure 3). A) Model fits by scaling different factors by latent patient estimates. BIC values shown relative to the scaling method providing the best fits. B) Cross-validation of model fits: relative log predictive probability of training fold model fits on held-out test folds, means across folds. For each fold, we used maximum likelihood estimation to obtain parameter fits across the four training folds. Those parameter fits were then used to generate predicted *P*(*Leave*) across bins in the held-out test fold patches, over which we summed log predictive probability of patch leaving decisions. C) Example schematics demonstrating how Models 2 (Left) and 3 (Right) make differing predictions for DV (Top) and *P*(*Leave*) (Bottom) on patches with rewards at *t* = [0,2] sec (‘R0R’ patches, black) compared with patches with rewards at *t* = [0,1] sec (‘RR0’ patches, blue). D) Per subject mean Simulated PRT for R0R versus RR0 patches from Model 2 fits (Left), Model 3 fits (Middle), and empirical mice PRT (Right). Points are colored per mouse. Axes are log-scaled. There was also a significant effect of greater PRTs for ‘R0R’ versus ‘RR0’ trials for Model 3 resulting due to a selection bias over latent patience values. E) Left: Model 3 free parameter fits versus mean PRT across mice (Left, *X*_0_: p = 0.009, Ψ: p = 0.0011, *P*_*max*_ : p < 0.0001, *R*: p = 0.0022, *ω*: p = 0.5828, *λ*: p = 0.2270, linear regression with Bonferroni correction). Right: Correlation matrix of Model 3 free parameter fit values across mice (significant correlations indicated with asterisks, Bonferroni correction). F) R^2^ statistics for predicting single trial PRT per mouse with a latent-scaled model 3 using a single shared set of fit parameters across mice, versus model 3 fit separately per mouse but without latent scaling (p < 0.0001, Wilcoxon signed-rank test). G) Single trial predictions from models fit in ‘F’. Example mouse 80 is shown (Top). Mouse with median fit difference across models, mouse 68, additionally shown (Bottom) as representative example since mouse 80 was an outlier for this fit comparison. H) First 20 patches from Fig. 3K showing single trial mouse and Model 3 predicted PRTs with reward timings indicated per patch.

**Figure S4:**
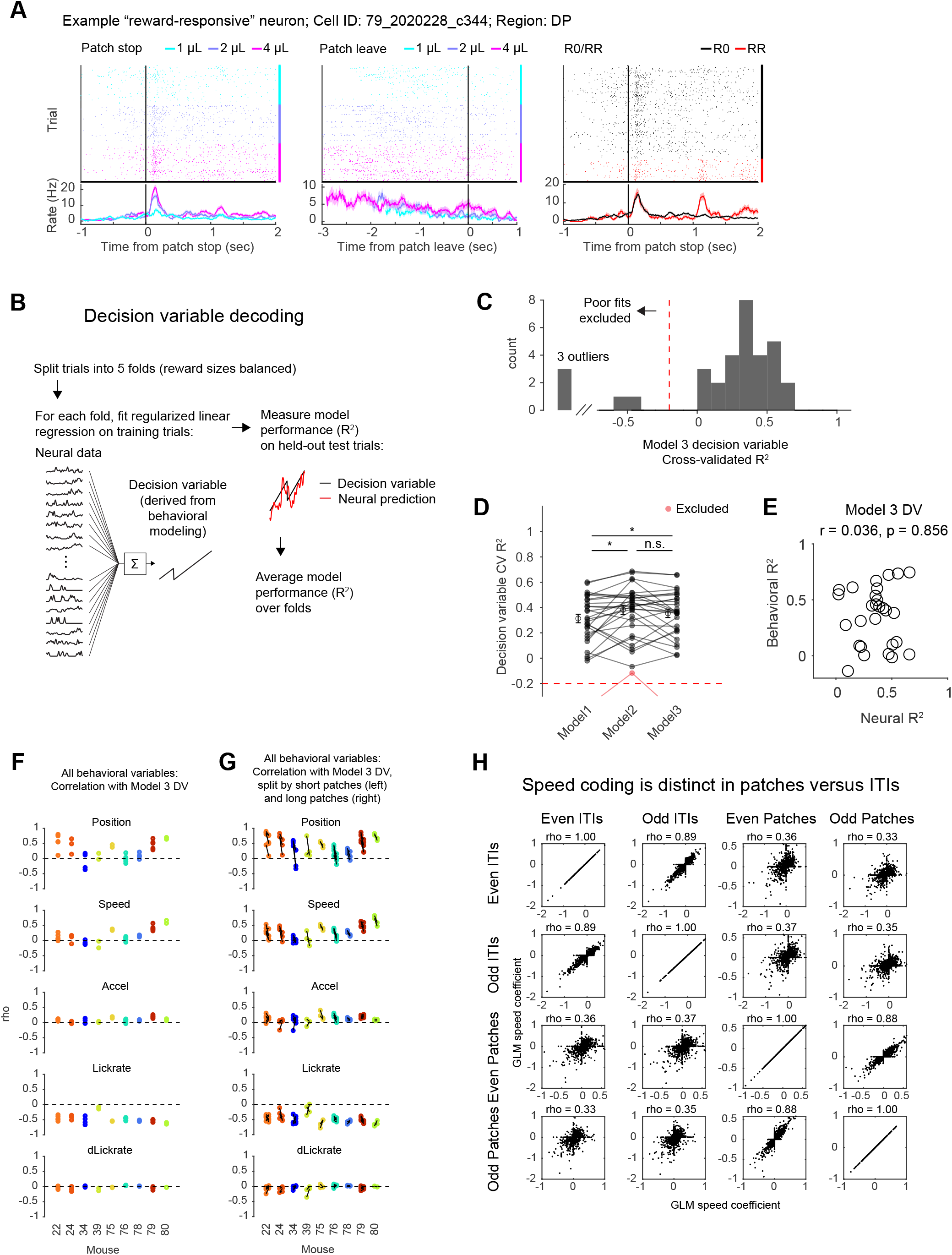
Additional example neuron, decision variable decoding, and behavioral confounds (related to Figure 4). A) An example neuron showing reward-responsive activity. Left: PSTH aligned to patch stop, split by reward size (cyan: 1 *μL*, purple: 2 *μL*, magenta: 4 *μL*; trials with reward at t=1 omitted). Middle: PSTH aligned to patch leave, split by reward size. Right: PSTH aligned to patch stop, split by whether reward was delivered at 1 second (red) or not (black); rewards of different size combined. B) Schematic of decision variable decoding (cross-validated linear regression of decision variables on smoothed, z-scored neural activity). C) Histogram of cross-validated *R*^2^ for each recording session in the data set. Sessions with CV *R*^2^ below an arbitrary threshold of −0.2 were deemed “poor fits” and excluded. D) Comparison of CV *R*^2^ for the decision variable derived from each of the models in Figure 3. * p < 0.05, Wilcoxon signed rank test (not corrected for multiple comparisons). E) *R*^2^ for Model 3 behavioral model fitting (y-axis) versus *R*^2^ for decoding of Model 3 decision variable from neural data (x-axis), after excluding 5 sessions with Neural *R*^2^ < −0.2. The ability of the Model 3 decision variable to predict behavior was not related to fidelity of neural decoding of that variable on a per-session basis. F) Pearson’s correlation between Model 3 DV (Fig. 3) and multiple behavioral variables: position, speed, acceleration, lick rate, derivative of lickrate. Each data point represents a single Neuropixels recording session. G) As in F, but split into short versus long patches (patch residence time above or below the median, per session). For each mouse, short patches are plotted on the left, and long patches on the right. Data points from the same session are connected by black lines. H) GLMs fit to intertrial intervals (ITIs) revealed qualitatively distinct speed coding compared to patches, suggesting that ramping activity in patches is not a low-level motor response. The structure of the GLM in patches was the same as in Figure 5, but fit to odd and even patches separately. The GLM in odd/even ITIs included the following regressors: Intercept, session time and its square, speed, and acceleration. Otherwise, the GLM fitting procedure in ITIs was identical to patches. In plots, each point represents a neuron, and “rho” above the plot indicates Pearson’s correlation coefficient.

**Figure S5:**
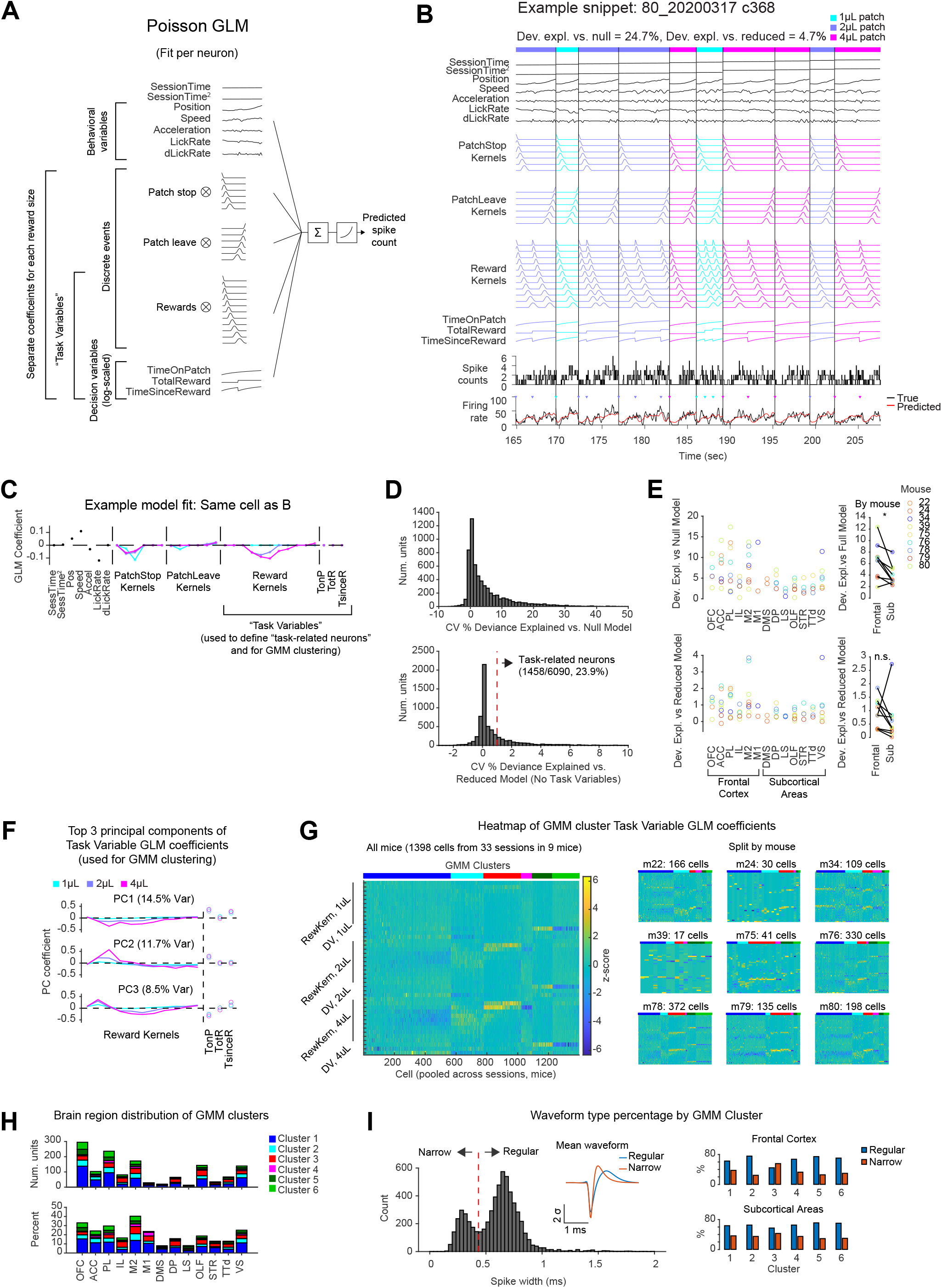
Additional information related to GLM fitting and GMM clustering (related to Figure 5). A) Schematic of the Poisson GLM. B) A snippet of an example cell showing the structure of the Poisson GLM: Regressors (top) and neural activity (bottom). Inverted triangles in the bottom panel indicate reward delivery. Vertical lines indicate the boundaries between patches; only on-patch times were included. Colors indicate reward size. Reward Kernels, Time on Patch, Total Reward, and Time Since Reward were considered “Task Variables.” C) GLM coefficients for the example cell shown in B. Task Variables were used for clustering (Reward Kernels, Time on Patch, Total Reward, and Time Since Reward; the latter three are also termed Decision Variables, or DVs). Only “task-related” neurons were included in clustering (panel D), and further were required to have at least one non-zero Task Variable GLM coefficient. D) **Top:** Cross-validated percent deviance explained versus null model (intercept only) for all neurons in the dataset. **Bottom:** Cross-validated percent deviance explained versus reduced model (no task variables) for all neurons in the dataset. Neurons with >1% deviance explained versus reduced model were designated as “task-related.” E) **Top:** Average percent deviance explained versus null model by brain region. Within each brain region, neurons were averaged by mouse. Colors indicate mice. **Bottom:** Average percent deviance explained versus reduced model by brain region. F) The top 3 PCs of the matrix of Task Variable coefficients (accounting for 34.7% of variance). Task Variable coefficients were z-scored prior to PCA. G) **Left:** Heatmap of z-scored GLM Task Variable coefficients for all neurons included in the analysis, split by cluster and sorted within cluster by the projection onto the first principal component (shown in F). **Right:** Same as left, but split by mouse. H) **Top panel:** Number of neurons in each GMM cluster, split by brain region. **Bottom panel:** Percent of all recorded neurons in each brain region assigned to each cluster. I) **Left:** Histogram of spike widths for all units in the dataset. A threshold of 0.433 ms was used to define Narrow versus Regular waveforms. Inset: The mean waveform for Regular and Narrow units. **Right:** Unit waveforms by GMM cluster. Especially in Frontal Cortex, Cluster 3 was enriched for units with Narrow waveforms. 56% of Frontal Cortex Cluster 3 neurons were narrow spiking, versus 32% of Frontal Cortex neurons from the other 5 clusters. In Subcortical Areas, these numbers were 43% versus 34%.

**Figure S6:**
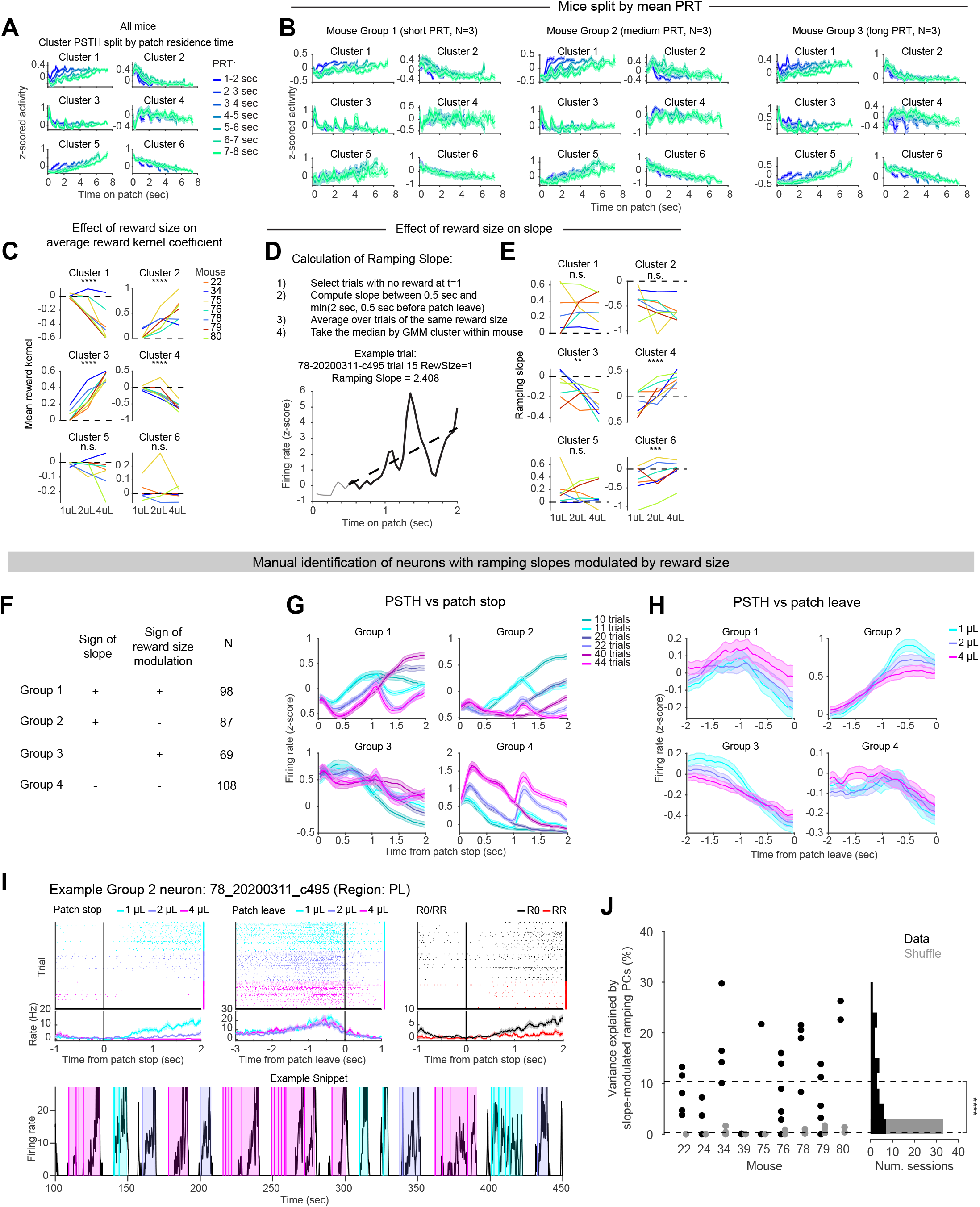
Additional analysis of ramping neural activity (related to Figure 6). A) Same as Figure 6A, but for all GMM clusters. B) Same as A, but with mice split into three groups by mean patch residence time (short, medium, long; N=3 mice in each group). Ramp-to-threshold-like activity in Cluster 1 could be observed in all three groups. C) Same as the top of Figure 6B, but for all GMM clusters. D) Explanation of how ramping slope is calculated, using trials with no reward delivery at t=1. E) Same as the top of Figure 6C, but for all GMM clusters. F) Neurons with ramping slopes modulated by reward size were identified using a shuffle analysis (see Methods). These neurons were then split into 4 groups based on the sign of the ramping slope and the sign of the modulation by reward size. G) PSTHs of z-scored neural activity aligned to patch stop for each of the 4 groups in G. Colors indicate patch types as in Figure 5E (“11 trials” indicates trials with 1 *μL* reward delivered at t=0 and t=1, etc.). Lines indicate means and shaded area indicates standard error of the mean. H) PSTHs of z-scored neural activity aligned to patch stop for the 4 groups in G, split by reward size. Lines indicate means and shaded area indicates standard error of the mean. I) An example Group 2 neuron showing ramping activity whose slope decreased with reward size. **Top:** Left: PSTH aligned to patch stop, split by reward size. Middle: PSTH aligned to patch leave, split by reward size. Right: PSTH aligned to patch stop, split by whether reward was delivered at 1 second (red) or not (black); rewards of different size combined. **Bottom:** A snippet of the same neuron’s activity across several consecutive trials, showing single neuron ramping activity over several seconds in individual trials. Shaded regions indicate patches, with the color indicating reward size. Vertical lines indicate reward delivery. J) Variance explained by principal components (PCs) of neural activity with ramping slopes modulated by reward size. Ramping PCs with reward size-modulated slopes were identified in the same manner as single units in panels G-J. Plot is analogous to Fig. 4E, but for ramping PCs with reward size-modulated slopes instead of all ramping PCs. Black points/bars indicate data, gray points/bars indicate a shuffle control in which PCs were circularly permuted relative to task events, once per session. **** P < 0.0001, data versus shuffle, sign rank test (N = 33 sessions).

## METHODS

### KEY RESOURCES TABLE

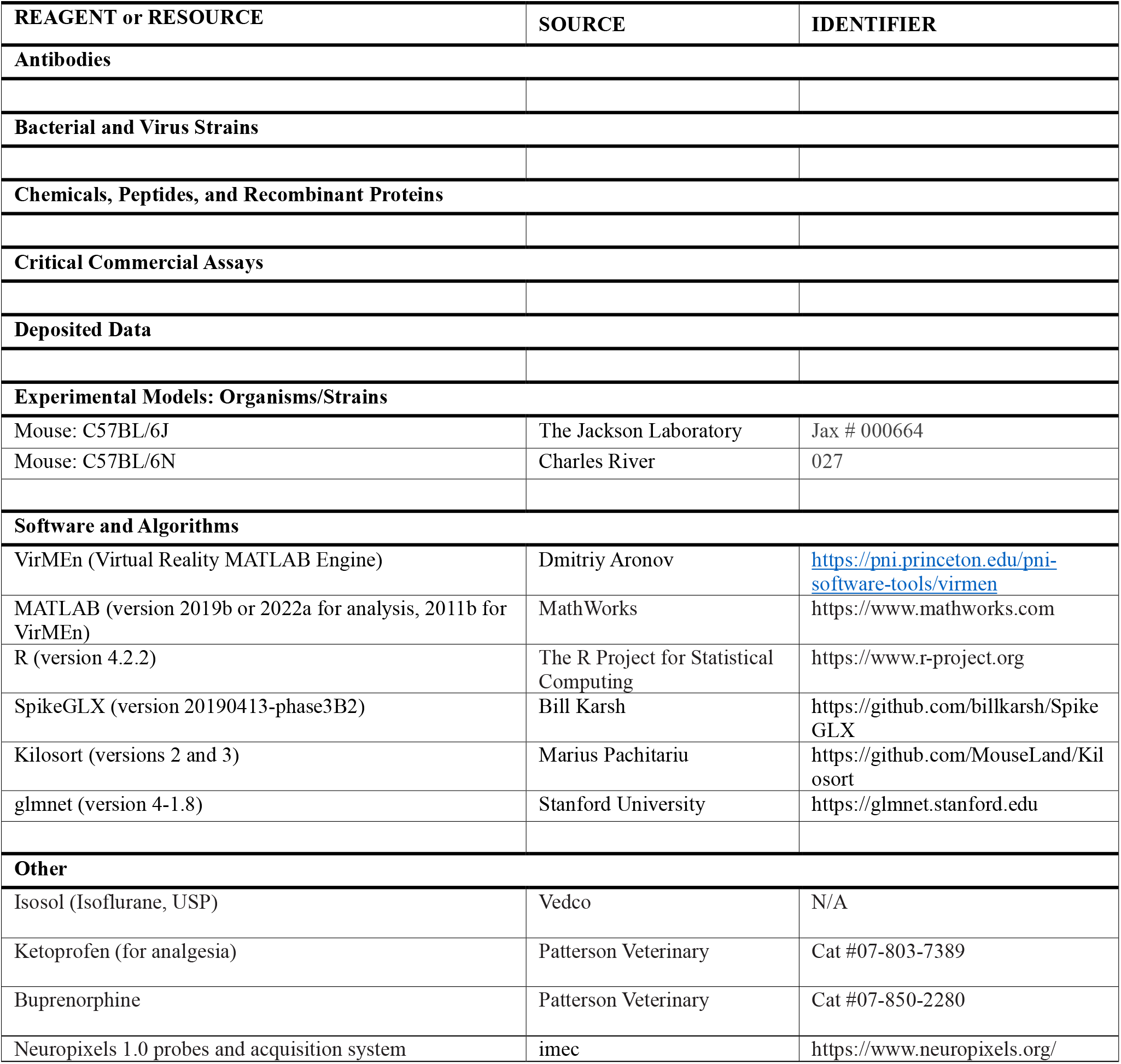

### LEAD CONTACT AND MATERIALS AVAILABILITY

Further information and requests for resources and reagents should be directed to and will be fulfilled by the Lead Contact, Naoshige Uchida. This study did not generate new unique reagents.

### EXPERIMENTAL MODEL AND SUBJECT DETAILS

#### Mice

A total of 30 adult mice (C57/BL6j, 20 male, 10 female, age 2-5 months) were used in the experiments. Mice were housed on a 12 h dark/12 h light cycle (dark from 07:00 to 19:00) and performed the task in the dark period. All procedures were performed in accordance with the National Institutes of Health Guide for the Care and Use of Laboratory Animals and approved by the Harvard Animal Care and Use Committee.

## METHOD DETAILS

### Surgery

Mice were anaesthetized using isoflurane (4% induction, 1-2% maintenance). For behavioral experiments, a custom titanium headplate was attached to the skull with Metabond (Parkell). For Neuropixels recordings, fiducial marks were made at the target sites for probe insertion using a fine-tipped pen, and a ground pin was inserted into the skull above contralateral cortex and attached with Metabond.

### Virtual reality setup

Virtual reality setups were identical to those used in ^73^. Three monitors (width 53 cm, height 30 cm) were placed in front and on either side of the animal. Virtual reality scenes were generated using VirMEn software in MATLAB^74^ on a workstation computer (DELL Precision 5810). Mice were head-fixed on a cylindrical Styrofoam treadmill (diameter 20.3 cm, width 10.4 cm). The rotational velocity of the treadmill was recorded using a rotary encoder. The output pulses of the encoder were converted into continuous velocity signal using custom Arduino code running on a microprocessor (Teensy 3.2). Velocity was integrated within VirMEn to compute position. Water was delivered to the mouse from a spout placed in front of the mouse’s mouth. Licks were monitored using an infrared sensor (OPB819Z, TT electronics). Voltage signals from the rotary encoder were digitized and recorded on the virtual reality computer using a data acquisition system (PCIe-6323, National Instruments). Water timing and amount was controlled using a solenoid valve (LHDA 1221111H, The Lee Company) and a switch (2N7000, On Semiconductor), with TTL pulses generated by the virtual reality computer via the PCIe-6323 data acquisition card.

### Patch foraging paradigm in virtual reality

Mice were water-restricted to maintain body weight >85% of their baseline weight to ensure task motivation while maintaining health. Additional water supplementation was provided shortly following training sessions with a total of task and post-task water ranging from 1.0-2.4mL per day, determined individually per mouse. Mice that showed task motivation at higher weights were given greater amounts of total water supplementation accordingly to minimize any stress from water restriction. Body weights ranged from 85-105% of baseline pre-restriction weight across mice.

#### Early training

Mice were habituated to the treadmill prior to initial head-fixation, delivering occasional water droplets from the reward spout while mice were free to explore on top of the treadmill, which was held in place to prevent rotation. After mice showed consistent interest over 1-2 sessions in consuming water droplets from the spout (2-3 days of habituation), head-fixation was begun in the following session. Initial head-fixed sessions were run for durations of 5-10 minutes, with gradual increases in session duration of typically 5-10 minutes across consecutive sessions, until reaching a max session duration of 45-75 minutes, varied per mouse based on how long they were observed to show task engagement as evident by running and licking behavior. Mice were initially head-fixed for 2-3 consecutive daily sessions before being given the following day off, after which the next set of consecutive daily sessions was increased by +1-2 days, followed by another day off, and continued in that manner until reaching a typical training schedule of 6 days on, 1 day off.

Initial head-fixed sessions were run in the dark. During those sessions, water rewards were delivered within 60sec intervals regardless of whether mice ran on the treadmill, to maintain positive reinforcement to rig fixation. To reinforce running behavior, mice were able to receive additional water rewards more readily via running, with water rewards delivered after traversing distances that were gradually increased across rewards.

After mice demonstrated consistent running behavior during the dark training (typically 2-3 sessions), mice began their initial training on the linear virtual track. After running a fixed distance, a visual “proximity cue” was illuminated on the track, followed by an automatic 1, 2, or 4µL (randomly varied) water reward delivery after a delay of 0.25-1.0s following cue onset. Running distance required to trigger the proximity cue and subsequent reward was increased after each block of 20 trials was completed. When mice completed >40 trials in a training session, the following session was started with a greater required running distance. This training stage served to teach mice that they must run to obtain rewards and that the proximity cue indicated an upcoming reward delivery. We determined that mice had successfully learned the association between the proximity cue and reward after observing consistent behavioral signs of reward expectation in response to proximity cue onset by reducing their running speed and licking. After mice progressed through increasing levels of running distance across sessions and demonstrated anticipatory slowdown and licking in response to the proximity cue onset prior to reward delivery, mice were transferred to the full version of the foraging task.

#### Full task

Mice traversed a virtual linear track wherein the periodic appearance of a visual proximity cue indicated that they could halt their running at that time to enter a virtual patch, in which water rewards could be obtained. Visuals for the linear track and proximity cues were the same as in earlier training. Each trial began with mice running along the track (travel time), and after running a fixed distance (∼2m; for two mice [34 and 39], the VR gain was set such that travel distance was ∼60 cm, with all other distances scaled accordingly), a proximity cue was illuminated. Contrary to earlier training, no automatic reward delivery occurred following the onset of the proximity cue. Instead, mice were required to pause their running on the treadmill while the cue was displayed, maintaining a near stationary positioning for ∼480ms, to obtain water reward(s). When mice achieved this stopping criteria, the surrounding visuals changed to indicate they had entered a patch, at which time an initial water reward was always delivered, droplet size = 1, 2, or 4µL, depending on patch type. If mice failed to reduce their speed sufficiently to meet the stopping criteria and continued running through the proximity cue (distance =∼40cm), the cue was extinguished and a new trial started at the beginning of the track.

Nine patch types were pseudorandomly varied across trials. Visual cues were the same across all trials, irrespective of patch type. Patches differed in their reward statistics, with size and frequency of reward deliveries varied across patch types. Three values were used for each, yielding nine unique combinations for patch type. Reward sizes were 1, 2, or 4µL, with size held constant across all rewards within a given patch. After the initial deterministic reward that was always delivered upon patch entry, additional reward events were drawn probabilistically following each one-second interval that elapsed on the patch. Values used for determining the probability of each reward delivery were dictated by an exponential decay function, which was scaled by reward frequency per patch type, *N*_0_ = .125, .25, *or* .5.

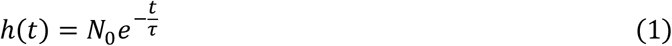

Values used for *N*_0_ per patch frequency type were .125, .25, or .5. A time constant of 8 seconds was used across all patches, *τ* = 8*s*. To obtain a discrete probability for reward delivery following each one-second interval, we integrate over the decay function:

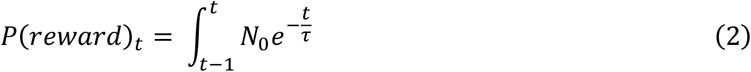

Mice were free to leave each patch at any time. Patches were delineated in virtual space, and leaving was triggered by crossing the virtual boundary (∼20cm from patch entry location). Leaving the patch marked the start of a new trial at the beginning of the track.

### Electrophysiological recording and spike sorting

Spiking data were collected using Neuropixels 1.0 single shank probes^75^. Neuropixels data were recorded with SpikeGLX (https://billkarsh.github.io/SpikeGLX/). Craniotomies were performed the day before recording and covered with Kwik-Cast (World Precision Instruments), and mice were allowed to recover overnight. On the day of recording, a 3d-printed piece was placed over the head of the mouse, blocking from the mouse’s sight the experimenter manipulating the probe above the mouse’s head. This was critical to maintain good behavioral performance on Neuropixels recording days.

Neuropixels probes were lowered using a Thorlabs micromanipulator (PT1-Z8) at 9 *μm*/sec.

After reaching the target depth, the probe was allowed to settle for 30 minutes prior to starting the recording. Recordings lasted 34 minutes (minimum) to 74 minutes (maximum) and were terminated once the experimenter determined the mouse was no longer engaging with the task (e.g., running through patches without stopping).

Neuropixels data were spike sorted offline with Kilosort 2 or 3^76,77^, followed by manual curation in Phy (https://github.com/cortex-lab/phy).

### Histology

After the last Neuropixels recording session, mice were perfused with phosphate buffered saline (PBS) followed by 4% paraformaldehyde in PBS. The brains were cut in 100 *μm* coronal sections using a vibratome (Leica). Brain sections were mounted on glass slides and stained with 4’, 6-diamidino-2-phenylindole (DAPI, Vectashield). Slides were imaged with a fluorescence microscope (Zeiss Axio Scan.Z1). Probe tracks were identified in histology images and aligned to a reference atlas using SHARP-Track^78^ or software written by Dr. Andrew Peters (https://github.com/petersaj/AP_histology).

## QUANTIFICATION AND STATISTICAL ANALYSIS

Unless otherwise noted, data were analyzed with custom code written in MATLAB (version 2019b or 2022a).

### Mouse inclusion criteria

A total of 30 mice were trained for these experiments, 22 of which (73.3%) passed our inclusion criteria for analysis. For inclusion, mice were required to progress through the early training stage to the full task and complete at least 10 sessions with at least 20 trials in each. Sessions were included after mice achieved >50% of their maximal session reward rate. 2 mice failed to progress through the initial training, and 3 mice failed to complete the required number of eligible sessions after progressing to the full task.

Of the remaining 25 mice that completed the required number of sessions, 3 were excluded from the analysis group due to their PRT varying randomly across patch types, indicating an indifference to reward statistics.

### Definition of latent state

To obtain an approximate measure of mice’s latent levels of patience across trials, we calculated a weighted average of PRT over successive trials. For each session, we ordered PRT by trial number. We then iterated over trials to compute the latent value per trial. First, we removed the PRT value associated with that trial, to prevent biasing the latent estimation. We then smoothed PRT across the session by applying a Gaussian filter (standard deviation = 10 trials, filter length = 30 trials). Artificial trials were added prior to the first trial of the session and following the last trial, using mean PRT from the first and last ten trials in the array, respectively, to avoid the filter being cutoff. After obtaining the smoothed PRTs across trials, we calculated the mean of the trials before and after the trial of interest and used that value as its latent patience estimate.

### Regression analyses modeling PRT

We fit a generalized linear mixed-effects model (GLME) with a log link function to predict PRT per patch type across subjects (Fig. 1G), given by the following equation:

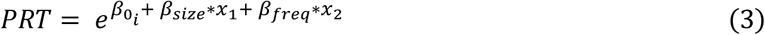

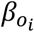 serves as a random effect, fit with a separate value per mouse (*i*). *β*_*size*_ and *β*_*freq*_ are fixed effects, each with a single parameter shared across the population. *x*1 and *x*2 indicate reward size and frequency, respectively, for each patch. The values for the three reward sizes and frequencies were standardized to [1, 2, *and* 4] to allow for interpretation of size and frequency coefficients on the same scale. The fit was performed using the ‘fitglme’ function in MATLAB version R2022a with the Statistics and Machine Learning Toolbox. The model was fit using the Laplace approximation, and the optimization was performed with ‘fminsearch’.

Additional GLME fits (same equation) were performed using 5-fold cross-validation and compared with alternative fits in which the trial labels (reward size, reward frequency, reward size and frequency, or mouse ID) were shuffled in the training set (Fig. S1H). Trials were split into folds by ordering patches per subject and labeling them 1-5 in serial. The GLME was then fit, as above, on the training folds. For the cross-validation and shuffles, GLME-predicted PRTs were calculated for patches in the held-out test folds.

We similarly fit a GLM using a log link function to predict PRT across patches by removing the per subject random effects term and adding a term to scale each patch by its latent ‘patience’ estimate (Fig. 2D), given by the following equation:

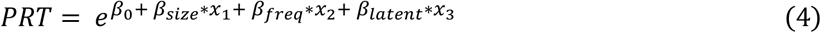

In this formulation, *β*_0_ is the log-intercept. *x*1 and *x*2 are the standardized values for reward size and frequency for each patch, same as in the GLME. *x*3 is the log of the latent ‘patience’ estimate for each patch. A single set of values for the four free parameters was fit across the population. We next performed this same GLM fitting procedure, but fitting a separate set of parameters individually for each mouse (Fig. S2F). We also performed several linear regressions, separately per mouse (Fig. S2A,E), in which the values for patch reward size and frequency were standardized the same as above.

### The Marginal Value Theorem (MVT)

In this study, we tested various predictions of the MVT. Following the derivation of the MVT by Charnov (1976)^3^ and Stephens and Krebs (1986)^1^, here we describe the assumptions and predictions of the MVT. We consider *n* patch types (*n* = 1,2,3, …, *n*). A patch type *i* is defined by *r*_*i*_(*t*), the instantaneous reward rate as a function of the time on the patch, *t*. We assume that the animal encounters each patch type with the probability *λ*_*i*_. We define *R*_*i*_ (*t*_*i*_) as the cumulative energy intake from staying in a patch type *i* for the duration *t*, where 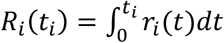. We also consider *E*_*S,i*_, which is the energy cost per unit time while searching in a patch type *i*. The gain function *g*_*i*_ (*t*) in a patch type *i* can be obtained by:

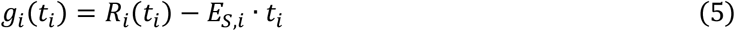

We assume that the derivative of *g*_*i*_(*t*_*i*_) is a monotonically decreasing function over time for all patch types.

We consider the time (*T*_*T*_) and energy cost per unit time (*E*_*T*_) of traveling between patches. The average time per trial to stay in a patch type (average travel time + average patch residence time) *T*_*u*_ is defined by:

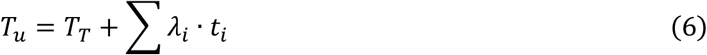

The average energy intake from a patch *E*_*e*_ is obtained by:

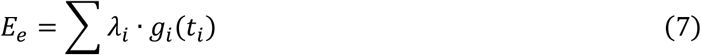

The overall rate of energy intake (*E*_*n*_) (energy intake on patch + energy loss while traveling) is then described by:

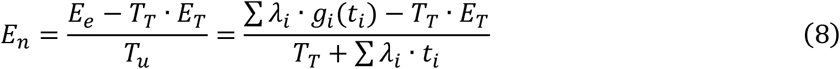

Assuming that the travel time is independent of the time spent in a patch, the above equation can be written from the standpoint of a patch type of interest *j* as:

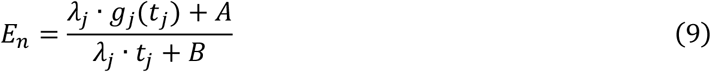

where *A* and *B* are not functions of *t*_*j*_.

To obtain a set of *t*_*j*_’s that maximizes *E*_*n*_, we differentiate *E*_*n*_ with respect to a given *t*_*j*_ (= patch residence time at patch *j*):

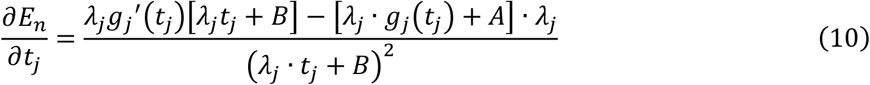

By setting 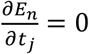, we obtain:

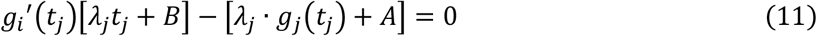

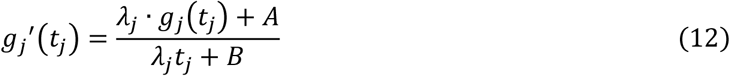

The right hand-side of this equation is equal to the overall rate of energy intake *E*_*n*_. Considering that *g*_*j*_′(*t*_*j*_) is a monotonically decreasing function, this indicates that an optimal forager should leave a given patch *i*, when the instantaneous energy intake rate *g*_*j*_′(*t*_*j*_) drops to the net rate of energy intake *E*_*n*_. This is Charnov’s Marginal Value Theorem (MVT).

Importantly, while *E*_*n*_, and therefore the patch residence time (*t*_1_, *t*_2_, ⋯, *t*_*j*_), depend on the values of travel time (*T*_*T*_), travel cost (*E*_*T*_), and search cost (*E*_*S,i*_), the following equality must hold true regardless of the values of these parameters:

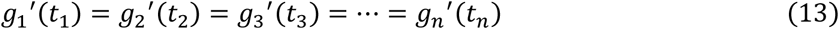

Note that *g*_*j*_*t*_*j*_ contains the cost of staying in the patch *j* (i.e. *E*_*S,j*_ · *t*_*j*_) in addition to the cumulative rewards *R*_*j*_(*t*_*j*_) as discussed above, with its derivative given by

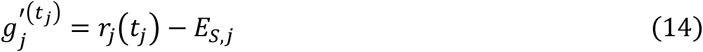

If the energy cost per unit time in a patch *E*_*S,j*_ is the same across all patches, like in our task, the following equality should hold true (**Prediction 1**):

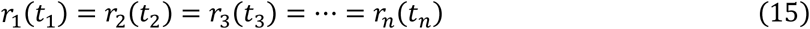

Next we consider the consequences of specific reward schedules used in our task such as exponentially-decreasing reward rates and the proportional scaling of overall reward rates between patches.

In our task, the instantaneous rate of reward for patch type *i* is:

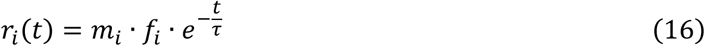

Where *μ*_*i*_ = [1,2,4], *f*_*i*_ = [0.125, 0.25, 0.5], and *τ* = 8 sec.

For a pair of two patch types, *i* and *j*,

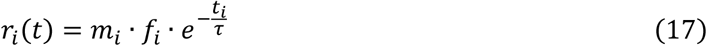

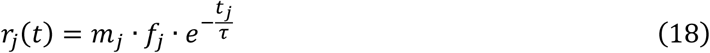

Now we define *a* such that 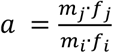.

At the MVT-optimal patch residence times, *t*_*i*_ and *t*_*j*_:

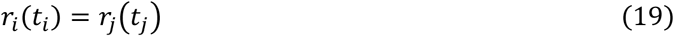

Therefore,

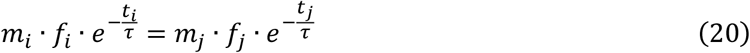

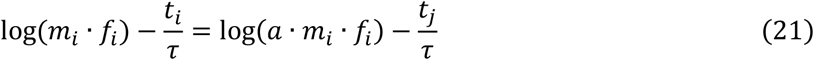

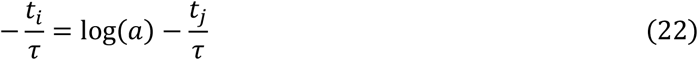

Thus,

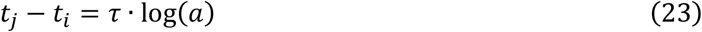

This means that although both *t*_*i*_ and *t*_*j*_ are sensitive to the net energy intake of the environment *E*_*n*_, the difference in patch residence times is constant for any given pair of patches with exponentially decaying reward rates with the same time constant (*τ*), irrespective of *E*_*n*_, as long as *a* is constant (**Prediction 2**).

Furthermore, with three patch types whose initial reward rates *m*_*i*_*f*_*i*_ increase proportionally such that:

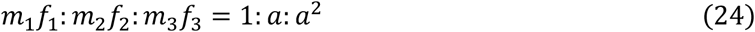

Then the difference in MVT-optimal PRT will be constant across successive patch types, i.e.,

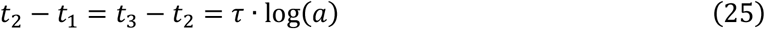

That is, the difference in the patch residence times should be constant across the pairs with the same proportional increase in reward rates (**Prediction 3**). In this study, we used patches with proportional reward sizes (1, 2, and 4μ*L*). The difference in leave time between 4 and 2 *μL* patches (*PRT*_4μ*L*_ − *PRT*_2μ*L*_) should, therefore, be equal to the difference between 2 and 1 *μL* patches, for patches with the same initial reward frequency. More specifically, this difference should be equal to a fixed number, 8 · log(2) = 5.5452 sec.

### Calculation of instantaneous expected reward rate

To determine whether mice are determining their patch leaving times via reward rate estimation and setting a threshold as MVT predicts, we calculate the instantaneous expected reward rate at the time of patch departure from an ideal observer perspective (same as Lottem et al.^54^). We use this quantification rather than the true empirical reward rate since the underlying reward frequency condition is a hidden feature of each patch and cannot be observed directly. The model assumes full knowledge of the distribution of patch types in the task and their respective reward statistics. Reward size is therefore always known, as the first reward on each patch provides this information. For each frequency condition, *f*_*i*_, likelihood is calculated based on the series of outcomes (rewards vs. omissions) observed across one second intervals within each patch o_1,_ …, o_n_. Assuming a uniform prior over frequency conditions, the probability of reward delivery at time *t*+1 is (same as Lottem et al.^54^):

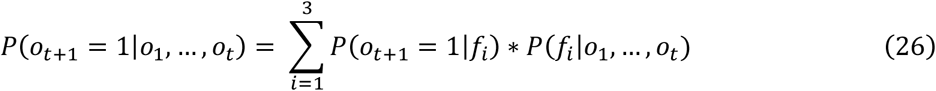

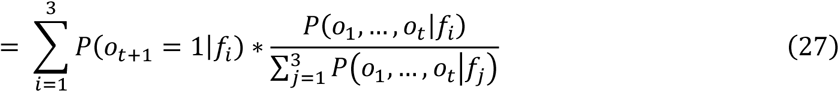

Probability of reward delivery at each timepoint per frequency condition is given by Eq. 2 and aggregate probability across observations per patch is:

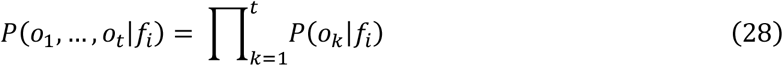

Lastly, to obtain instantaneous reward rate from expected reward probability, we scale it by reward size and consider that expected value over a one second interval as reward rate.

### Behavioral model fitting

Models 1 and 2 have five free parameters: *X*_0_ sets the midpoint of the sigmoid. *Ψ* is inverse temperature. *maxP*_0_ constrains the models’ *P*(*Leave*) output. *ω*_0_ is an exponent for power-law scaling of slope per reward size. *λ*_0_ is an exponent for power-law scaling by latent state. Model 3 included those five, plus one additional free parameter for the value subtracted from the integrator per reward delivery, *R*.

The slope of the integrator model was scaled based on the patch’s reward size. We fit this slope scaling by a power-law function to allow for varying sensitivity to reward size across mice. We used μ = [.5, 1, 2] for [1, 2, 4] μ*L*, respectively, to normalize slope to the medium reward size.

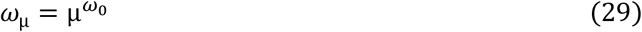

Slope was additionally scaled by the estimate of patience for each patch (patch = *i*). We again used a power-law function for this scaling, fitting the free parameter, *λ*_0_, per mouse to account for differences in how dependent PRT was on estimated patience across mice. Values for patience across mice were mean-normalized to ensure fit values across parameters could be straightforwardly compared across mice. Each patch’s normalized latent, *L*_*i*_, was scaled by *λ*_0_ to determine its contribution to slope scaling:

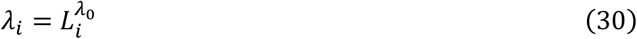

Because variance in patch leaving times increases when PRTs are longer, we also scaled *μLxP* with patches’ latent patience estimates. Because 0 < *maxP* < 1, we used a function that would reasonably allow scaling in either direction, while constraining the transformed value to remain between 0 and 1:

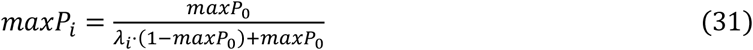

The other free parameters, *Ψ*, and, for Model 3, *R*, were not scaled across patches.

The models produce a decision variable (DV) per one second interval (*t*) by calculating its current integrator value, *X*_*i,t*_ and subtracting *X*_0_ from it:

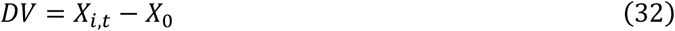

The models differ in how they each track time, TOP (Time on Patch) or TSLR (Time Since Last Reward), and whether they integrate rewards delivered (*nRews* = number of rewards delivered up to that point on the patch), to compute *X*_*i,t*_:

Model 1, reward indifferent:

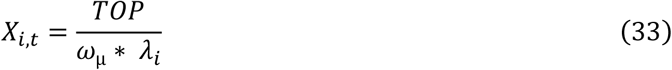

Model 2, reward reset:

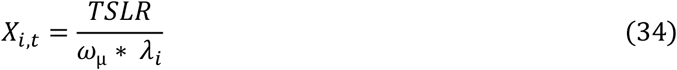

Model 3, reward integrator:

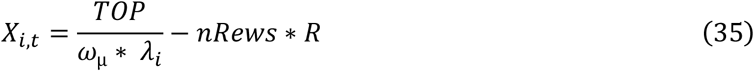

To compute the model’s output prediction, *P*(*Leave*), the probability of leaving a patch per one second bin, the DV is scaled by inverse temperature and sigmoid transformed with the output constrained by *maxP*:

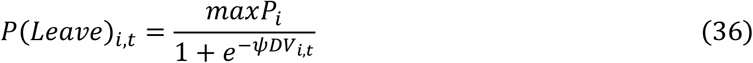

We optimized free parameter fits per subject and compared the models using the mfit toolbox for MATLAB. Maximum a posteriori estimates were computed using MATLAB’s fmincon function using 20 random initializations, converging upon the set of parameter fits that maximized log likelihood of the models at predicting *P*(*Leave*) across one second time bins. Ranges for the parameters used to fit were *X*_0_[−5: 20], *Ψ* [0: 10], *maxP*_0_[.01: .98], *ω*_0_ [0: 2], *λ*_0_[0: 4], and *R*[0: 20], with uniform priors for each. Resulting parameter fits are shown in Fig. S3. BIC values were computed for each model fit, and a protected exceedance probability was calculated for each model to estimate how prevalent the model is across subjects.

5-fold cross-validation was additionally performed across models. Trials were split into folds by ordering trials per subject and labeling them in serial 1-5. Fitting was then performed as above on the training folds, after which we computed the log predictive probability over trials from the held-out test folds.

### Using behavioral models to patch leaving times on single trials

To compute the average leaving time predicted from a given behavioral model on a single trial (Fig. 3L-N), we first calculated the “leave density” per each one-second time bin. For time bin in each patch, the leave density is given by the product of the fraction of trials the model predicted to still be on the patch at the start of the time bin by that time bin’s predicted leave probability: *Fraction on patch*_*t,i*_ * *P*_*t,i*_(*Leave*) = *Leave density*_*t,i*_. The predicted leaving time is then computed by averaging time bins, weighted by the leave density.

### Unit inclusion criteria

To be included for analysis, units from Neuropixels recordings had to have a minimum firing rate of 1 Hz within patches and additionally at least 1 Hz within each third of the session to ensure that they were tracked throughout the recording session.

### Spike waveforms

Spike waveforms were extracted for each unit using C_Waves (https://billkarsh.github.io/SpikeGLX/). Spike width was defined as the difference between the time of the waveform minimum and the maximum following this minimum. A threshold of 0.433 ms was used to define Regular (>0.433 ms) versus Narrow (<0.433 ms) waveforms (Fig. S5I).

### Principal components analysis (PCA) of neural activity

For each unit, firing rate traces were generated by binning spikes into 50 ms bins. These traces were then smoothed by convolution with a causal filter, a half-Gaussian, where the standard deviation of the full Gaussian was 100 ms. Prior to PCA, only in-patch times were selected, excluding the inter-patch intervals, and this trial-concatenated, smoothed firing rate trace of each neuron was z-scored. PCA was performed on the resulting per-session matrix of neurons x in-patch time points (MATLAB function “pca”).

### Definition of ramping principal components (PCs)

To identify “ramping” PCs (Fig. 4E), trials with no reward at t=1 were selected. For each PC, slopes were computed on single trials by calculating the linear slope between t=0.5 and t=2 or patch leave time, whichever came first. A sign rank test was performed to compare these single trial slopes against 0, and PCs were deemed “ramping” if this test was significant at p < 0.05.

### Linear models to predict behavioral decision variables from neural activity

For each unit, spikes were binned into 50 ms bins and smoothed with a 100 ms Gaussian kernel to generate single-cell firing rate traces within each patch. Then, firing rate traces were z-scored across patches for each neuron, so that model coefficients could be compared across neurons. We then fit cross-validated linear models to predict the ramping decision variable (DV) from neural activity.

Regularized linear models were fit using the function *lasso* in MATLAB with alpha = 0.1 (10% L1, 90% L2 regularization), predicting a DV from the activity of all recorded neurons at each time point. For cross-validation, patches were split into the same 5 folds as for the GLM (see above). The regularization parameter lambda was selected by nested cross validation within training folds.

Specifically, the training data was split into 10 folds and models were fit with a range of lambda values. Mean squared error was computed on nested test folds for each value of lambda, and lambda was chosen such that mean squared error was within 1 standard error of the minimum. The linear model was then fit on the entire training set to obtain *β*, the coefficients mapping neural activity to the DV, and *R*^2^ was computed on the test set. *R*^2^ and *β* were averaged across training folds to obtain final values for each recording session/neuron.

### Generalized linear model (GLM) to predict neural activity from behavioral variables

Poisson GLMs were fit to spiking activity of single neurons using the glmnet toolbox (https://glmnet.stanford.edu)^79,80^. To prepare data for GLM fitting, we binned spikes from each neuron into 50 ms bins aligned to patch stop and continuing until patch leave. The following variables were included as regressors in the GLM:

1. Session time: time in session, (time in session)^2^
2. Behavioral variables: patch position, velocity, acceleration, lick rate, d(lick rate)
3. Patch stop kernels: 6 kernels x 3 (1 set per reward size), peaks spanning 0-1 seconds from patch stop
4. Patch leave kernels: 6 kernels x 3 (1 set per reward size), peaks spanning 0-1 second prior to patch leave
5. Reward kernels: 11 kernels x 3 (1 set per reward size), peaks spanning 0-2 seconds after each reward
6. Time on patch: 3 (1 per reward size). Time on patch was log scaled: *t*_*scaled*_ = In(*t* + 1)
7. Time since last reward: 3 (1 per reward size). Time since reward was log scaled: *t*_*scaled*_ = In(*t* + 1)
8. Total reward: 3 (1 per reward size). Total reward was log scaled: *R*_*scaled*_ = In(*R* + 1)

Thus, a total of 85 variables were used in the GLM (86 including intercept). Kernels were a raised cosine basis with log scaling of the x-axis (time), created using MATLAB code from the Pillow lab (https://github.com/pillowlab/raisedCosineBasis). Parameters used were: logScaling = ‘log’, logOffset = 1.0, timeRange = [0 5].

Variables were z-scored prior to model fitting. Variables with one coefficient per reward size were z-scored across reward sizes, so that coefficients could be compared across reward sizes. Kernels were z-scored together.

Data were prepared for glmnet using custom MATLAB code, and then fit using glmnet (version 4.1-8) in R (version 4.2.2). Model fitting was performed on the Harvard Faculty of Arts and Sciences Researching Computing cluster (FASRC cluster, https://www.rc.fas.harvard.edu/).

Models were regularized with elastic net regularization with *α* = 0.9 (90% L1, 10% L2) to encourage sparse coefficients^81^. For cross-validation, patches were split into 5 folds. Each patch was assigned in its entirety to a single fold. Folds were counterbalanced for patches of different reward size. Cross-validation with these folds was used to select the regularization parameter for each neuron (using the function *cv*.*glmnet*).

Cross-validated deviance explained was computed for each neuron as follows:

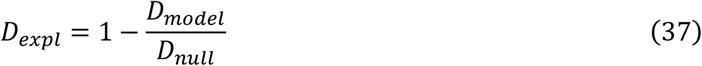

Where *D*_*model*_ is the cross-validated model deviance and *D*_*null*_ is the cross-validated null deviance. These two terms were computed once per test fold and then averaged across folds. For each test fold 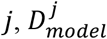 and 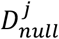 were computed using the formula for Poisson deviance:

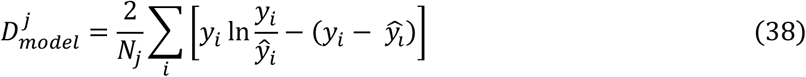

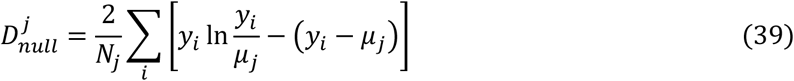

Where *N*_*j*_ is the number of observations (time bins) in the *j*th test fold, *y*_*i*_ is the number of spikes in the *i*th time bin in the *j*th test fold, *ŷ*_*i*_ is the predicted mean of the Poisson distribution in the *i*th time bin (computed using the model fit on the training data), and *μ*_*j*_ is the mean spike count per bin in the training data. For bins with no spikes, we set 0 ⋅ In(0) = 0.

To estimate the contribution of task variables (reward kernels, time on patch, total reward, and time since reward) to model fits, we fit a reduced model in which task variables were removed. Poisson deviance was computed for the reduced model and deviance explained by task variables was computed as 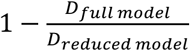. Neurons with deviance explained > 1% by this metric were deemed “task-related.”

### Unsupervised clustering of GLM coefficients

We used a Gaussian Mixture Model (GMM) to cluster the matrix of Task Variables (M=42) x Task-Related Neurons (N=1398; we excluded 60 neurons whose coefficients on task variables were all zero, leaving 1398 task-related neurons in this analysis) (Fig. 5). The dimensionality of this matrix was reduced to 3 × 1398 using PCA (34.7% of variance; Fig. S5F). The dimensionality-reduced matrix was clustered using a Gaussian Mixture Model (function fitgmdist in MATLAB, with parameters RegularizationValue = 0.4, MaxIter = 10000, Replicates = 10, TolFun = 10^−6^) with K (the number of clusters) spanning 1-10. For each value of *K*, the Bayes Information Criterion (BIC) was calculated. *KK* was selected to minimize BIC (K = 6; Fig. 5C).

### Identification of neurons and principal components with ramping slopes modulated by reward size (Fig. S6F-J)

We identified ramping neurons whose ramping slopes were modulated by reward size. Ramps could be positive or negative, and reward size modulation could also be positive or negative, resulting in four groups of neurons (Fig. S6F). For each neuron, single-trial ramping slopes were computed as in Fig. S6D. To be considered “ramping,” a neurons’ single trial ramping slopes had to differ significantly from zero (one-sample t-test p < 0.05). To assess whether a neuron’s single trial ramping slopes were “reward size-modulated,” first, single-trial ramping slopes were linearly regressed against the reward size on each trial; neurons with a positive regression coefficient were considered to have ramping slopes positively modulated by reward size, and vice versa. Then, reward size labels were shuffled across trials, 100 times per neuron, and a z-test was performed to compare the regression coefficients for the unshuffled versus shuffled data. Neurons with z-test p-value < 0.05 were considered to have significantly “reward size-modulated” ramping slopes. To produce the neuron grouping in Fig. S6F-I, neurons had to pass both preceding tests at p < 0.05 ([1] ramping slope different from zero across reward sizes, and [2] shuffle test for reward size-modulation), and the resulting neurons were split by whether the average ramping slope was positive or negative and whether reward size modulation was positive or negative, resulting in four groups (Fig. S6F).

To estimate the fraction of in-patch neural activity variance accounted for by ramping activity whose slope is modulated by reward size, we performed the same analysis on principal components of neural activity, and summed the variance explained by principal components identified as having reward size-modulated ramping slopes (Fig. S6J). For details on how principal components were computed, see the section “Principal components analysis (PCA) of neural activity.”

### State space modeling of ramping and stepping dynamics

The ramping model was implemented as a constrained recurrent, switching linear dynamical system with Poisson emissions^82,83^. The model has a continuous latent state *x* and discrete latent state *z* that both evolve over time. The discrete latent state can be in one of two states corresponding to an accumulation state (*z*_*t*_ = *acc*) and a boundary state (*z*_*t*_ = *b*). The continuous latent state is initialized to *x*_0_ < 1. In the accumulation state, it follows a drift-diffusion process with a separate slope per reward size and a shared diffusion variance *σ*^2^. It also accumulates pulsatile reward inputs *u*_*t*_ with a separate accumulator weight per reward size. This corresponds to the following dynamics

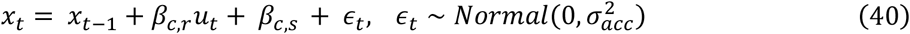

where *β*_*c,r*_ is the coefficient for rewards for reward size *c*, and *β*_*c,s*_ is the slope for reward size *c*. When *β*_*c,r*_ < 0 the reward decrements the accumulator, although importantly *β*_*c,r*_ is unconstrained during fitting and is initialized at 0. In the boundary state, the continuous dynamics have no constant slope and a small constant variance. However, because a salient reward event could alter an animal’s decision to leave even after that intention was formed, rewards still influence the continuous state which can cause the model to leave the boundary

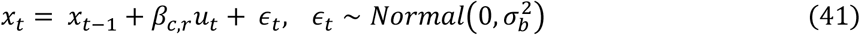

where 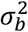 is fixed to 1e-4. Transitions between discrete states depends on the value of the continuous state relative to a fixed boundary threshold at *x* = 1. In particular, the discrete state transition distribution is

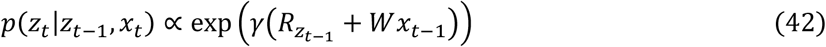

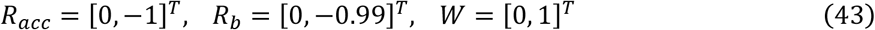

and *γ* = 50. This setting of the parameters places a boundary at *x* = 1 with probabilistic transitions to and from the boundary state for values of the continuous state near the boundary. We note that the discrete state is slightly encouraged to stay in the boundary state, as in the discrete state the threshold for returning to the accumulation state decreases to *x* = 0.99. Overall, the generative model is

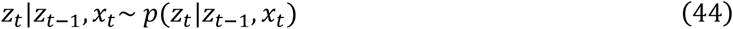

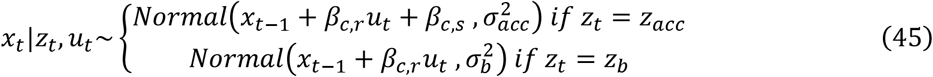

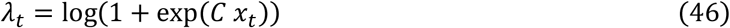

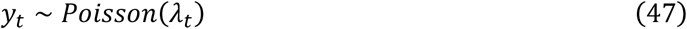

The time-varying reward inputs are shifted in time such that *u*_*t*_ is equal to one 400ms after a reward and otherwise equal zero.

The parameters 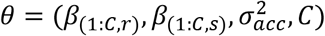 were fit using variational Laplace EM (vLEM)^83^, while the other parameters were fixed to correspond to the ramping model structure. The vLEM algorithm returns an approximate posterior distribution *q*(*x*_1:*t*_) that approximates the true posterior distribution over the continuous latent states *p*(*x*_1:*T*_ ∣ *y*_1:*T*_, *u*_1:*T*_, *θ*). For model comparison, the held-out log-likelihood was estimated via importance sampling using *q*(*x*_1:*t*_)

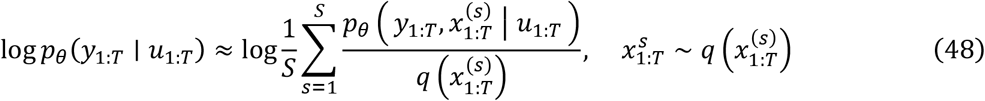

with S = 200 and where the discrete latent state is marginalized in the numerator when computing *p*_*θ*_.

The stepping model was implemented as a constrained HMM with input-dependent transitions and trial-dependent rates. It has two discrete states corresponding to a “down” and “up” state. Here the time-varying inputs *u*_*t*_ are one-dimensional and equal to 1 when there is a reward at time *t* and are otherwise zero. As in the ramping model, the rewards are shifted 400ms in time. The model starts in the down state and in the absence of a reward transitions to the up state with probability *p*_*s*_. When there is a reward, the model is forced to transition back to the down state. The generative model is

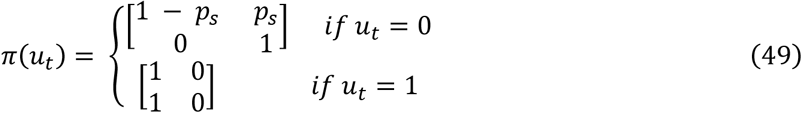

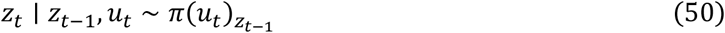

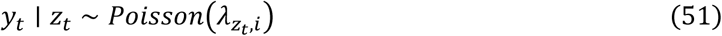

where 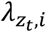 is the firing rate for state *z*_*t*_ on trial *i*. Notably, we fit separate firing rate parameters for each state and trial to allow the model to account for variability in overall rates across trials. While the ramping model has condition-dependent parameters and the stepping model does not, the stepping model gains significant flexibility by having per-trial firing rate parameters.

The overall model parameters are *θ* = (*λ*_:,1:*I*_, *p*_*i*_) and are fit via maximum likelihood using the EM algorithm. The per-trial firing rates are initialized to the mean number of spike counts in the first and last 10 time bins of a trial for the down and up states, respectively. To evaluate the model on heldout trials, the step probability *p*_*s*_ was kept constant but we fit the per-trial firing rates 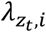 to the test trials. Given the per-trial firing rates on test trials, the heldout log-likelihood was computed as the log-likelihood of the test trials marginalizing out the discrete states. This is a strong baseline since it has access to the heldout data for fitting the per-trial firing rates, whereas the ramping model does not have per-trial parameters that are fit on the heldout data.

For both models, we fit the model to the summed spike count of Cluster 1 neurons from the frontal cortex. We fit and compared the models for sessions with at least 10 such neurons. The models were fit to binned spike counts in 50ms bins. For each session, 80% of the trials were used for training the model parameters and the held-out loglikelihood is computed and reported on the remaining 20% of the trials. Both models were fit using code in the SSM (https://github.com/lindermanlab/ssm) and SSM-DM (https://github.com/davidzoltowski/ssmdm) code packages.

## Acknowledgements

We thank Ed Soucy and Brett Graham of the Harvard Center for Brain Science Neurotechnology Core for engineering assistance, Emmanuel Garrison-Hooks for assistance with mouse training, and members of the Uchida lab for valuable discussions and comments on the manuscript. We thank Dr. Matteo Carandini, Dr. Kenneth Harris, Dr. Andrew Peters and other members of the Cortex lab for their advice on Neuropixels recording. This work was supported by grants from the NIH Brain Initiative (U19NS113201, to S.W.L. and N.U.), the Simons Foundation (SCGB 697092, to S.W.L. and N.U.), the Sloan Foundation (S.W.L.), the McKnight Foundation (S.W.L.), NIH T32MH020017 (M.B.), NIH F32MH126505 (M.G.C.), and the Wu Tsai Neurosciences Institute (D.Z.).

## Author Contributions

M.B. and N.U. designed the foraging task. M.B. built the behavioral and recording setups with assistance from H.K. M.B., M.G.C., L.K., and J.S. collected behavioral data. M.B., M.G.C, and J.S. analyzed behavior with feedback from N.U. M.B. and M.S.T. constructed the integrator models with guidance from J.D. M.B., M.G.C., and L.K. collected Neuropixels data. M.G.C., M.B., and J.S. analyzed Neuropixels data with feedback from N.U. D.Z. performed the recurrent linear dynamical systems modeling under the supervision of S.W.L. M.B., M.G.C. and N.U. wrote the manuscript. All authors reviewed the manuscript.

## Data and Code Availability

Data and code will be posted to online repositories upon publication.

